# Zasp52 strengthens whole embryo tissue integrity through supracellular actomyosin networks

**DOI:** 10.1101/2022.10.11.511783

**Authors:** Dina J. Ashour, Clinton H. Durney, Vicente J. Planelles-Herrero, Tim J. Stevens, James J. Feng, Katja Röper

## Abstract

During morphogenesis, large scale changes of tissue primordia occur that are all coordinated across an embryo. In *Drosophila*, several tissue primordia and embryonic regions are bordered or encircled by supracellular actomyosin cables, junctional actomyosin enrichments networked between many neighbouring cells. We show that the single *Drosophila* Alp/Enigma family protein Zasp52, which is most prominently found in Z-discs of muscles, is a component of many supracellular actomyosin structures during embryogenesis, including the ventral midline and the boundary of the salivary gland placode. Zasp52, we uncover, contains within its central coiled-coil region a type of actin-binding motif usually found in CapZbeta proteins, and this domain displays actin binding activity. Using endogenously-tagged lines we identify that Zasp52 interacts with a subset of junctional components, including APC2, Polychaetoid/ZO-1, and Sidekick, as well as actomyosin regulators. Analysis of *zasp52* mutant embryos revealed that the severity of the embryonic defects observed scales inversely with the amount of functional protein left. Large tissue deformations occur at sites where actomyosin cables are found during embryogenesis, suggesting a model whereby supracellular cables containing Zasp52 aid to insulate morphogenetic changes from one another. This is further supported by the analysis of *in vivo* processes as well as by an *in silico* vertex model of morphogenetic domains separated by a stiffness boundary.

## Introduction

Morphogenesis, the generation of shape in development, is the basis of organ formation and major topological changes in development. Within individual cells changes in shape are driven by the cytoskeleton. Over the last decade, enormous progress has been made in deciphering which type of cell shape changes drive changes at the tissue scale (Blanchard et al., 2009; Guirao et al., 2015; Herrera-Perez and Kasza, 2018). Cell shape changes in most cases depend on the actomyosin cytoskeleton underlying the plasma membrane, the actomyosin cortex. This cortex is linked to cell surface receptors such as cadherins or integrins, thereby allowing modulation of the actomyosin cortex during cell-shape changes through cell-cell and cell-matrix adhesion (Chugh and Paluch, 2018; Mege and Ishiyama, 2017).

Crucially, during tissue morphogenesis cells do not change shape in isolation, rather a close coordination between all cells in a tissue is essential for productive changes at the tissue scale. Such coordination is based on different molecular mechanisms. Many morphogenetic events in embryos occur in epithelial tissues, where cell-cell adhesion between neighbouring cells allows for physical coupling of behaviours. The degree of turnover and stability of adhesion sites between neighbouring cells determines the extent of physical coupling between the neighbours, thereby for instance controlling the amount of neighbour exchanges (Song et al., 2013). Mechanical coupling through cell-cell adhesion connects the actomyosin cortices of neighbouring cells, which can also tie into cytoskeletal structures such as actin stress fibres or the microtubule cytoskeleton. Within epithelial cells, actomyosin is enriched near apical adherens junctions in what has been historically described as the ‘actin belt’ or ‘adhesion belt’ (Furukawa et al., 2017). Interestingly, this cell-cell-adhesion-associated actomyosin can be coordinated between neighbouring cells into seemingly “supracellular” assemblies. These were first observed during wound healing in embryonic epithelia (Martin and Lewis, 1992), termed actomyosin ‘cables’ or ‘purse strings’, and subsequently found during morphogenetic processes across the evolutionary tree (Rodriguez-Diaz et al., 2008). Actomyosin cables have been intensively studied in model processes such as dorsal closure and salivary gland invagination in the *Drosophila* embryo (Jacinto et al., 2002; Röper, 2012; Sidor et al., 2020), neurulation in both *Ciona* (Hashimoto and Munro, 2019) and mouse (Galea et al., 2017) and wound healing (Wood et al., 2002). Actomyosin assemblies can show different properties and provide different functionalities: they can be stable, or dynamic and short lived, be involved in active morphogenetic changes or stable delineation of differently fated populations of cells (Röper, 2013). In a few instances, the upstream molecular cascade leading to the assembly of the cable has been elucidated (Hashimoto and Munro, 2019; Röper, 2012; Sidor *et al*., 2020).

We have previously analysed the morphogenesis of the tubes of the salivary glands from a flat epithelial primordium in the *Drosophila* embryo termed the salivary gland placodes (Suppl.Fig. S1 illustrates the tube morphogenesis of the salivary glands), as well as the mechanism that leads to the assembly of supracellular actomyosin cables surrounding each of these two bilateral placodes (Röper, 2012; Sanchez-Corrales et al., 2021; Sanchez-Corrales et al., 2018; Sidor *et al*., 2020). The salivary gland placodes are specified on the ventral side of the embryo half-way through embryogenesis at late stage 10, and cells start to invaginate through a focal point termed the invagination pit in the dorsal-posterior corner, forming a narrow lumen tube on the inside whilst cells continuously invaginate (Girdler and Röper, 2014; Sidor and Röper, 2016). Cells at the invagination point undergo apical constriction driven by apical-medial actomyosin prior to internalisation, whilst cells further away from the invagination pit undergo directional cell intercalations driven by polarised junctional actomyosin, to continuously feed cells towards the invagination point (Sanchez-Corrales *et al*., 2021; Sanchez-Corrales *et al*., 2018). Furthermore, the assembly of the supracellular actomyosin cable at the boundary of each placode with the surrounding epidermis is dependent on the anisotropic localisation of the transmembrane protein Crumbs and its downstream effectors, Pak1 and aPKC (Röper, 2012; Sidor *et al*., 2020). The cable is under increased tension compared to nearby junctions that are not part of the cable, as shown by laser-ablation of junctions that form part of the cable and measurements of initial recoil of junction vertices (Röper, 2012). The function of the circumferential cable surrounding the placode is not yet clear, but it could help the cell internalisation by exerting an inward-directed boundary force, or it could help to insulate cell behaviours within the primordium from the surrounding epidermis.

Zasp52 is a member of the Alp/Enigma family of proteins, containing an N-terminal PDZ domain and four LIM domains (Fig. 1A). *Drosophila* Zasp52 was originally identified as a component of striated muscles, localised to the Z-line of these, where it interacts with its binding partner α-actinin and is important to link actin-barbed ends into the Z-disc (Jani and Schock, 2007). More recently, Zasp52 was also found to localise to the actomyosin cable that assembles at the leading front of epidermal cells during dorsal closure in the fly embryos (Stronach, 2014). These cells move towards the dorsal side of the embryo to cover the non-embryonic tissue of the amnioserosa. Zasp52 was shown to assist in providing a straight leading-edge front during dorsal closure, thereby assisting the correct matching of segments between left and right sides of the embryo (Ducuing and Vincent, 2016).

**Figure 1.**
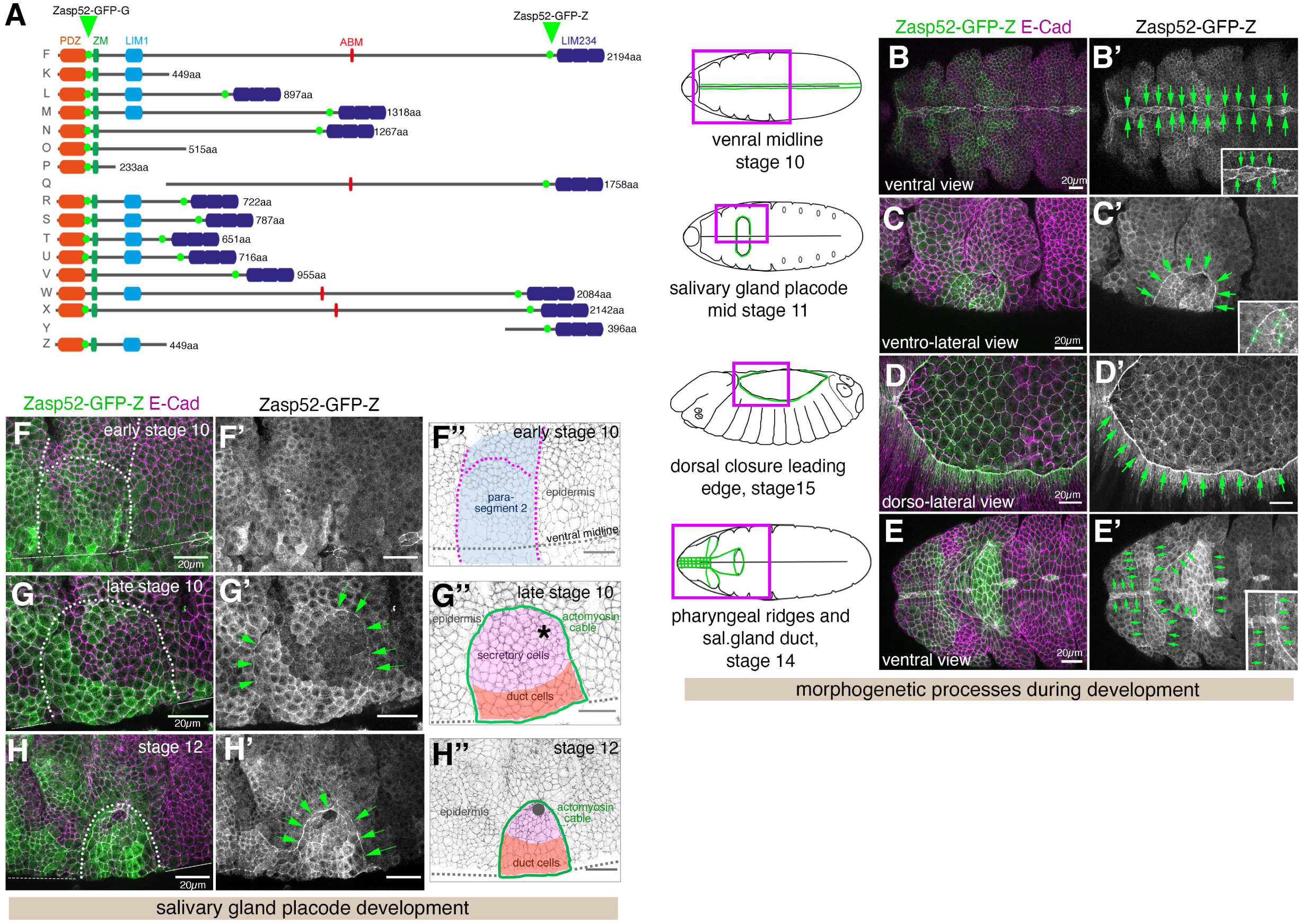
Localisation of endogenous Zasp52 to actomyosin cables in the Drosophila embryo. **A** Predicted protein isoforms of Zasp52 likely expressed in the *Drosophila* embryo, length in amino acids is given. The full-length protein contains an N-terminal PDZ domain, followed by a Zasp motif (ZM), a first LIM domain (LIM1), a long unstructured region of various length containing the newly identified actin-binding motif (ABM), followed by three C-terminal LIM domains (LIM 2-4). The insertion sites of two GFP-protein traps (line Zasp52-GFP [G00189], Zasp52-GFP-G, and Zasp52-GFP [ZCL423], Zasp52-GFP-Z, are indicated by green arrow heads and dots. **B-E’’** Localisation of Zasp52-GFP (Zasp52-GFP [ZCL423]) to various supracellular actomyosin cables in the *Drosophila* embryo: the cables lining the ventral midline (**B**, **B’**), the cable surrounding the salivary gland placode (**C**, **C’**), the leading edge epidermal cable during dorsal closure (**D**, **D’**) as well as other ventral actomyosin cables during head involution surrounding the pharyngeal ridges and cells forming the salivary gland duct (**E**, **E’**). Magenta boxes in the embryo schematics indicate positions of images taken, green lines indicate the actomyosin cables and assemblies shown in the images. Green arrows in **B’-E’** indicate the position of the actomyosin cables that Zasp52 localises to. Insets show higher magnifications of sections of the images. Circles in the inset in **C’** highlight stronger accumulation of Zasp52 in tricellular junctions. **F-H’’** Within the salivary gland placode, Zasp52-GFP-Z initially localises in a patchy pattern during specification of the tissue in parasegment 2 (**F**-**F’’**), before starting to accumulate at the boundary cable from late stage 10 on (**G**-**G’’**), before more strongly accumulating there over time (**H**-**H’’**). **F’’-H’’** show schematics of the stages, cell types (secretory cells, pink, versus duct cells, orange) and position of the forming invagination pit (asterisk in **G’’**) as well as of the actomyosin cable at the boundary (green). Green arrows in **G’** and **H’** point to Zasp52 accumulation at the position of the cable. Zasp52-GFP-Z is in green in **B-H** and as a single channel in **B’-H’**, E-Cadherin to label cell outlines is in magenta in **B-H**. Scale bars are 20µm. See also Supplemental Figures S1 and S2.

Here, we show that Zasp52 is in fact a component of many, though not all, embryonic actomyosin cables or supracellular actomyosin assemblies. Zygotic loss of Zasp52 leads to a reduction of junctional F-actin in the salivary gland placodal cable. We identify a novel F-actin binding motif (ABM) in the coiled-coil domain of Zasp52 that is related to the motif usually found the in the actin-capping protein CapZbeta. In addition, we identify several ubiquitous cell-cell-adhesion associated proteins as interaction partners of Zasp52, in particular APC2, Polychaetoid and Sidekick (Ahmed et al., 2002; Choi et al., 2011; Finegan et al., 2019; Letizia et al., 2019). Embryos lacking all of maternal and zygotic Zasp52 function show major defects in morphogenetic events in tissues all associated with the presence of supracellular actomyosin cables such as the salivary glands, the ventral midline, the head and the leading edge/amnioserosa interface. We propose that the supracellular actomyosin cables containing Zasp52 serve as mechanical insulators, preventing morphogenetic changes from ‘spilling over’ into neighbouring regions or otherwise interfering with nearby events, a model which is further supported by *in silico* investigations.

## Results

### Zasp52 is a component of embryonic supracellular actomyosin cables

In order to analyse Zasp52 protein localisation in the *Drosophila* embryo we made use of two protein-trap insertion lines in the *zasp52* locus (Fig. 1A). Zasp52-GFP (using either the Zasp52[ZCL423] or Zasp52[G00189] lines, with the former shown here) within the embryonic epidermis of *Drosophila* was particularly strongly localised to junctions that were part of supracellular actomyosin assemblies or cables (Suppl.Fig. S1 F-H). Both GFP-exon trap lines label most isoforms of Zasp52, including the longest possible one (Fig. 1A, protein isoform F). Zasp52-GFP was enriched in junctions of the ventral midline (Fig. 1 B, B’), the cable surrounding the salivary gland placode (Fig. 1 C, C’), the leading edge of epidermal cells during dorsal closure (Fig. 1 D, D’) as well as several large-scale cables that can be found on the ventral anterior side of the embryo at the start of head involution (Fig. 1 E, E’; (Röper, 2013)). These ventral anterior cables in particular appeared linked into a large-scale network that spans the whole anterior epidermis at this stage (arrows in Fig. 1E’). We could not detect Zasp52-GFP though in cables found at parasegmental boundaries in the early embryo before stage 11 or in the more dynamic and short cables found in tracheal placodes during tracheal morphogenesis (Suppl.Fig. S2). In addition to supracellular cables, Zasp52-GFP localised at lower levels to some bicellular junctions, and was always slightly more enriched at tricellular junctions (highlighted in Fig. 1C’ inset).

We analysed in more detail the localisation and levels of Zasp52-GFP in the forming salivary gland placode. Zasp52-GFP was only present at low levels in the early embryonic epidermis at early stage 10 prior to placode specification (Fig. 1F-F’’), but showed a patchy expression in the early salivary gland placode at late stage 10, with accumulation at circumferential cable commencing at this stage (Fig.1 G-G’’). During stage 11 and 12 Zasp52-GFP quickly accumulated more and more in the forming actomyosin cable at the placode boundary at (Fig. 1H-H’’).

Zasp52 therefore appears to be an enriched component of many, though not all, supracellular actomyosin cables in the *Drosophila* embryo.

### Zasp52 contains an actin-binding motif related to the CapZbeta-actin binding motif

The strong enrichment of Zasp52-GFP in many supracellular actomyosin structures and especially the circumferential cable around the salivary gland placode prompted us to investigate whether Zasp52 played a role in the establishment or function of this cable. In wild-type embryos, not only myosin II (Suppl.Fig. S1 F-F’’) but also F-actin accumulates strongly at the boundary of the placode (Fig. 2 A-A’’ and (Röper, 2012)). In embryos zygotically lacking Zasp52, using a deletion of *zasp52* called *zasp52Δ* (Jani and Schock, 2007), this F-actin accumulation at the placode boundary was strongly reduced, as it was also in transheterozygous embryos of the *zasp52Δ* allele combined with an allele of a deficiency spanning the *zasp52* locus (Fig. 2B and C).

**Figure 2.**
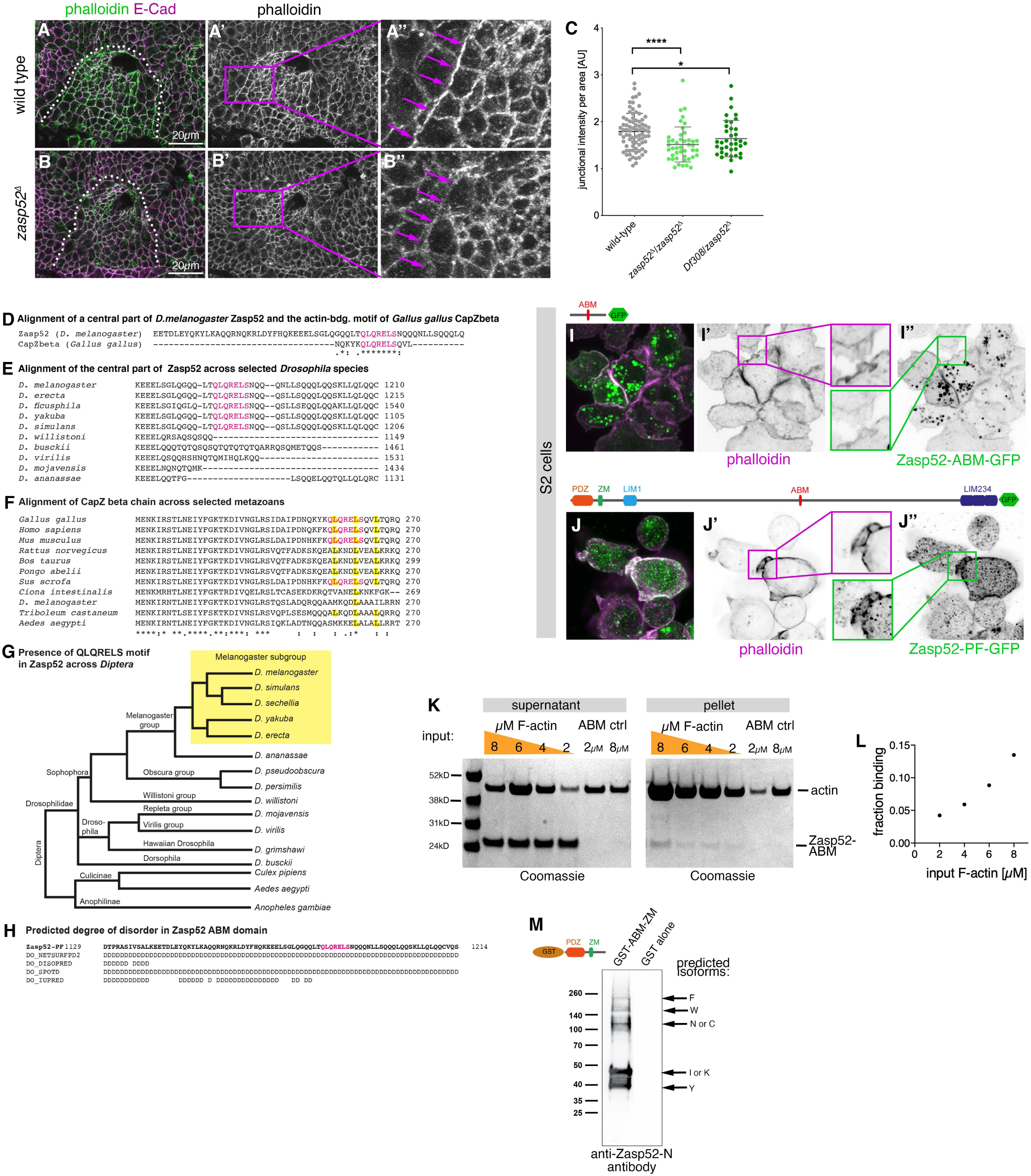
Zasp52 contains a novel F-actin binding site related to CapZbeta and affects supracellular F-actin localisation. **A-C** F-actin accumulates at the boundary of the salivary gland placode in wild-type embryos where a supracellular actomyosin cable forms (**A**-**A’’**). In embryos zygotically mutant for *zasp52*, a deletion mutant termed *zasp52^Δ^*, less F-actin accumulates (**B**-**B’’**). E-Cadherin to label cell outlines is shown in magenta and phalloidin to label F-actin in green. **A’’** and **B’’** are magnifications of the areas highlighted in **A’** and **B’**. **C** Quantification of phalloidin labelling intensity at the boundary of the placode in the wild type compared to either *zasp52^Δ^* homozygous mutant embryos or embryos carrying one copy of *zasp52^Δ^* and one copy of a deficiency uncovering the *zasp52* locus, *Df 308*. Statistical significance was determined using two-tailed Mann-Whitney test, shown are mean +/- SD; * <0.05; **** <0.0001. **D** Pairwise alignment of CapZbeta (*Gallus gallus*) and Zasp52 (*Drosophila melanogaster*) with the conserved motif highlighted in pink. Note that the motif identified in the central region of Zasp52 is near identical to the actin-capping motif in CapZbeta. **E** Amino acid sequence alignment of Zasp52-PF across members of the *Drosophila* genus. Conserved, putative actin binding motif highlighted in pink. **F** Amino acid sequence alignment of CapZbeta across metazoan species with conserved leucines reported to mediate binding of the terminal helix of CapZbeta highlighted in yellow (Hug *et al*., 1992; Kim *et al*., 2010) and the complete QLQRELS motif (pink letters). **G** Phylogenetic tree of the *Drosophila* genus with highlighted melanogaster subgroup (yellow) in which species show the QLQRELS motif in Zasp52. In **D** and **F** an (* asterisk) indicates positions which have a single, fully conserved residue, (: colon) indicates conservation between groups of strongly similar properties, (. period) indicates conservation between groups of weakly similar properties. **H** Secondary structure predictions about local disorder in the stretch of amino acids surrounding the ABM in Zasp52-PF, generated with Quick2D toolkit (NeetSurfP2 (Klausen et al., 2019) ; DISOPRED3 (Jones and Cozzetto, 2015); SPOT-Disorder (Hanson et al., 2017)) suggesting almost the entire region of the protein to be disordered (D = disorder). **I-I’’** Ectopic expression of the Zasp52-ABM fused to GFP in S2 cells. Zasp52-ABM-GFP (**I’**’, green) localises to the cell cortex where it colocalises with F-actin labelled by phalloidin (**I’**, magenta). Zasp52-ABM-GFP also shows unspecific aggregates in the centre of the expressing cells. **J-J’’** Ectopic expression of the long Zasp52 PF isoform fused to GFP in S2 cells. Zasp52-PF-GFP (**J’**’, green) localises to the cell cortex where it colocalises with F-actin labelled by phalloidin (**J’**, magenta). Zasp52-PF-GFP also shows small unspecific aggregates in the centre of the expressing cells. **K** Ultracentrifugation-based co-sedimentation assay of F-actin and Zasp52-ABM. 0.8µM Zasp52-ABM was incubated with increasing concentrations of pre-polymerised F-actin (2µM, 4µM, 6µM, 8µM). Blots of supernatant and pellet were analysed by staining with instant blue dye. 2µM and 8µM Zasp52-ABM were centrifuged without F-actin as control. **L** Fraction of bound Zasp52-ABM plotted against F-actin concentration. **M** Pull-down of bound Zasp52 isoforms using GST-Zasp52-PDZ-ZM as bait, revealed by labelling with an anti-Zasp52-N antibody, indicating that this domain likely participates in dimerisation of Zasp52. See also Supplemental Figure S3.

Zasp52 has been suggested to interact with F-actin directly through its N-terminal region, containing the PDZ domain and Zasp motif (Liao et al., 2020). Using a purified Zasp52 fragment containing the PDZ and Zasp motif (PDZ + ZM) we confirmed *in vitro* that this binding, although weak, is direct and does not require additional factors (Suppl. Fig. 3). Furthermore, in the Z-disk of muscles Zasp52’s binding to actin is mediated by its binding partner α-actinin (Jani and Schock, 2007). However, using a GFP protein-trap insertion into the α-actinin locus we found that α-actinin-GFP in embryos did not localise to junctions of epidermal cells and was not enriched in the cable surrounding the salivary gland placode (data not shown). As actin-binding through the PDZ + ZM part of Zasp52 appeared weak, we analysed the amino acid sequence of the longest possible isoform of Zasp52 (isoform PF) in more detail for further possible conserved actin-binding motifs that might have been overlooked. We identified a motif in the central unstructured region of Zasp52 that closely resembles the highly conserved actin-capping motif usually found in the actin-capping protein CapZbeta (Fig. 2 D-G, (Hug et al., 1992; Kim et al., 2010)). In CapZbeta, the conserved residues are part of a flexible C-terminal domain that has been shown to be able to flip in and out of the hydrophobic furrow of the F-actin analogue Arp1-A. This interaction is integral to the capping of the barbed end of filamentous actin (Narita et al., 2006; Urnavicius et al., 2015; Yamashita et al., 2003). While the motif in Zasp52 is not located at the C-terminus, it is located in an area of higher disorder implying increased flexibility (Fig. 2H). This motif in Zasp52 is not only found in *D.melanogaster*, but conserved in a subset of *Drosophilidae* (Fig. 2G).

We expressed the Zasp52-actin binding motif (ABM; see location in Fig. 1A)) fused to GFP in S2 cells, where it colocalised with cortical F-actin structures labelled using phalloidin (Fig. 2 I-I’’). Furthermore, the Zasp52-PF isoform, containing the ABM in its central part, also colocalised with F-actin labelled by phalloidin in S2 cells (Fig. 2 J-J’’). We then purified recombinant Zasp52-ABM and performed *in vitro* actin-binding assays. Zasp52-ABM pelleted with F-actin in an F-actin-dose dependent manner (Fig. 2 K-L).

Additionally, we uncovered that a GST-tagged N-terminal fragment of Zasp52 comprising the PDZ domain and Zasp motif (ZM) can co-precipitate several untagged endogenous Zasp52 isoforms from embryonic lysate (Fig. 2M), supporting that this domain is involved in Zasp52 dimerisation or multimerisation, which has been suggested to be dependent on the interaction of the Zasp motif with the LIM1 domain (Gonzalez-Morales et al., 2019).

Thus, Zasp52 is important for F-actin accumulation in the cable surrounding the salivary gland placode, possibly via the newly-identified actin-binding motif in its central region. The ability to dimerise or multimerise suggests that Zasp52 might not only bind and cap actin filaments with the two respective actin-binding motifs, but that it could also assist in actin cross-linking, actin bundling or other higher order organisation of F-actin filaments, thereby possibly affecting actomyosin contractility at the site of supracellular actomyosin structures.

### Zasp52 associates with junctional proteins

In order to identify if and how Zasp52 was recruited specifically to supracellular actomyosin structures and uncover what function it might fulfill there, we set out to identify potential further interaction partners in addition to F-actin. To do so, we generated lysate from embryos prior to formation of muscles (pre stage 14), either from embryos expressing endogenously tagged Zasp52, *Zasp52[ZCL423]*, or from wild-type embryos, or from embryos expressing an endogenously tagged Armadillo/μ-Catenin, *armadillo-YFP*, and performed anti-GFP co-immunoprecipitations (note that as soluble cold lysates were used, no F-actin or polymerized microtubules are present in the input material). Anti-GFP immunoprecipitation of Zasp52-GFP lysates compared to wild-type lysates showed a specific enrichment of many potential interactors with links to both actomyosin and cell-cell adhesion (Fig. 3 A; i.e. APC2, Sidekick, Polychaetoid, Patj, Raskol, Canoe, Cindr, Bazooka, p120catenin, alpha-Catenin, G-beta13F, Scribble), whereas the anti-GFP immunoprecipitation from Armadillo-YFP lysates compared to wild-type lysates identified the components of adherens junctions known to interact with it as enriched (Fig. 3B; i.e. Armadillo, alpha-Catenin, Shotgun, APC2, p120catenin,). The identification of in part distinct sets of interactors for Zasp52-GFP and Armadillo-YFP indicated that Zasp52 is not just a direct interactor of adherens junctions (Fig. 3A-C; for a full list of interactors see Suppl. Tables 1 and 2).

**Figure 3.**
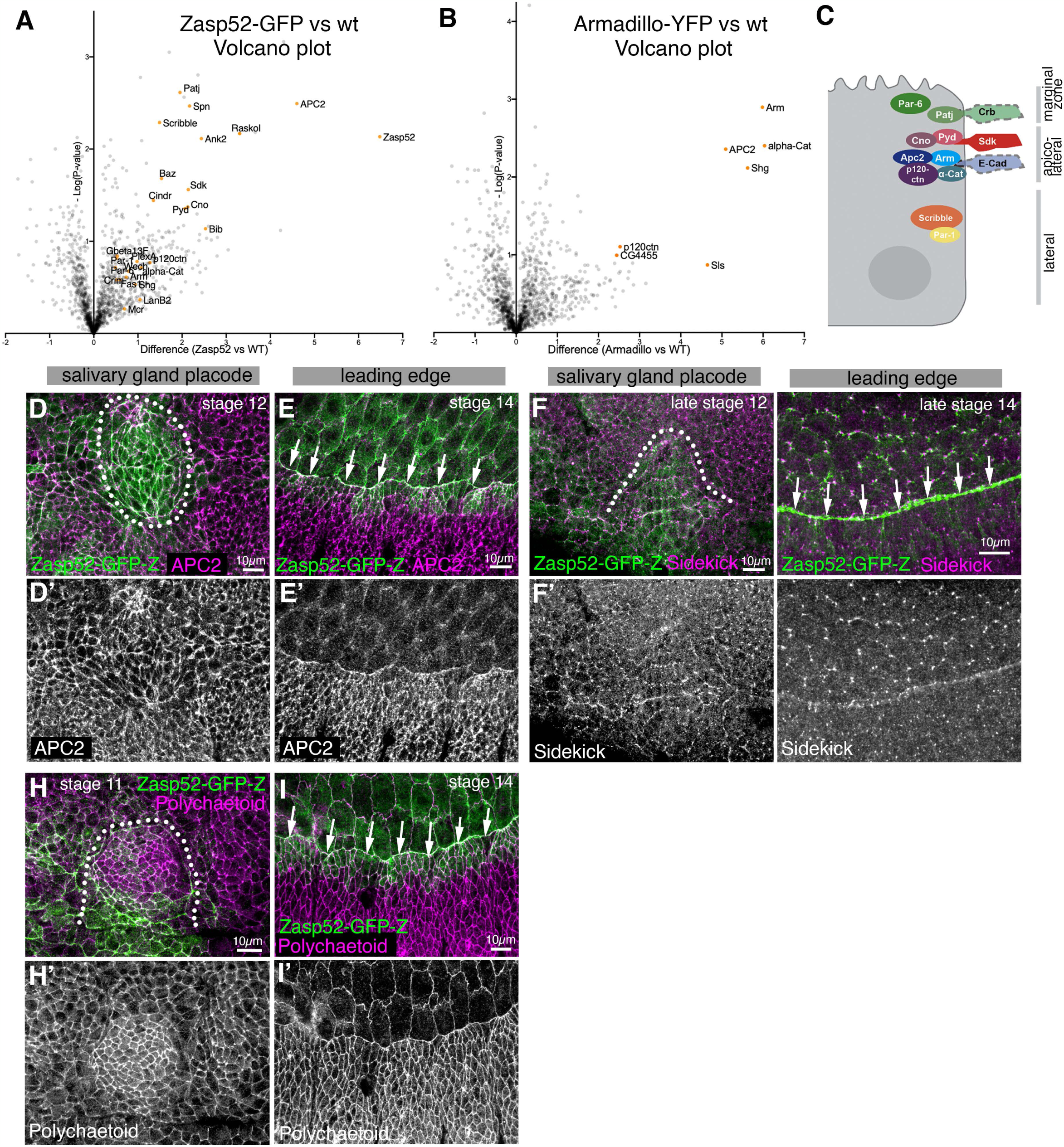
Zasp52-GFP interacts with many junctional proteins. **A, B** Volcano plots comparing spectral intensity values generated from anti-GFP co-immunoprecipitations from embryo lysates of Zasp52-GFP, Armadillo-YFP and wild-type embryos. Three experiment each for Zasp52-GFP and wild-type embryos were performed and two for Armadillo-YFP embryos. **A** shows enriched interactors for Zasp52-GFP compared to wild type, and **B** shows enriched interactors for Armadillo-YFP compared to wild-type. In **A**, hits related to cell adhesion and regulation of junctional cytoskeleton are labelled, and in **B** the top interactors that include the well-known Armadillo interactors at junctions. x-axis is the difference score for each protein as calculated in MaxQuant, y-axis is the probability score calculated between Zasp52-GFP or Armadillo-YFP and wild-type. The significance threshold was set to 0.75. **C** Schematic of different protein complexes and interactions in marginal zone, apico-lateral and lateral junctions in *Drosophila* epithelial cells. **D-I’** Localisation of three possible Zasp52-interaction partners within the embryonic epidermis. Localisation in and around the salivary gland placode is shown, as well as localisation in the leading edge epidermal cells as well as the amnioserosa during dorsal closure, two processes where a prominent supracellular actomyosin cable forms that Zasp52 localises to. **D-E’** APC2 localises to apical junctions, both bi-cellular and tri-cellular, but is more enriched at tri-cellular junctions. It is not enriched in either actomyosin cable. **F-G’** Sidekick is particularly enriched in tri-cellular junctions, but also in some bi-cellular ones. It is not enriched in either actomyosin cable. **H-I’** Polychaetoid/ZO-1 is localised homogeneously to apical junctions throughout the epidermis. It is not enriched in either actomyosin cable. Zasp52-GFP is always in green, APC2, Sidekick and Polychaetoid/ZO-1 in green or as individual channels. Dotted lines indicate the boundary of the salivary gland placode. Scale bars are 10µm. See also Supplemental Figure S4 and Supplemental Tables S1 and S2.

We analysed the localisation of potential interaction partners in comparison to Zasp52-GFP, in particular the localisation of the co-immunoprecipated and junctional components APC2, Polychaetoid/ZO-1 and Sidekick (Ahmed *et al*., 2002; Choi *et al*., 2011; Finegan *et al*., 2019; Letizia *et al*., 2019). APC2, one of the two APC tumour suppressor proteins in *Drosophila* with functions in Wnt signalling and cell adhesion (Ahmed *et al*., 2002; Hamada and Bienz, 2002), was localised to junctions in the embryonic epidermis and was particularly enriched at tricellular junctions, but was not specifically enriched in supracellular actomyosin structures (Fig. 3 D-E’, salivary gland placode and leading edge during dorsal closure are shown as examples). However, it colocalised with Zasp52-GFP in the junctions where Zasp52 was enriched. Similarly, Sidekick, a protein especially enriched in tricellular junctions in the embryonic epidermis and important for epithelial integrity during morphogenesis (Finegan *et al*., 2019; Letizia *et al*., 2019), also colocalised with Zasp52-GFP in these junctions (Fig. 3 F-G’). Lastly, Polychaetoid/ZO-1, a protein with a variety of functions in apical junctions including linkage to the actin cytoskeleton (Choi *et al*., 2011), localised to apical junctions in the embryonic epidermis and was enriched in placodal junctions, again colocalising there with Zasp52-GFP (Fig. 3 H-I’).

Thus, Zasp52-GFP appeared to associate with many junctional proteins at bi- and tri-cellular junctions. When analysing its localization along the apical to basal extent of the epithelial junctions, Zasp52 in actomyosin cables colocalised with E-Cadherin at the level of adherens junctions, but also partially overlapped with the more apically localised Crumbs protein within the apical marginal zone of the lateral sides (Fig. 3C and Suppl. Fig. S4 A-F).

The associated proteins analysed above do not usually show a specific enrichment in supracellular junctional actomyosin structures, suggesting that the localisation to and incorporation of Zasp52 into such structures is likely controlled by its pattern of expression and its general ability to bind F-actin and junctional components, rather than a unique membrane-associated binding partner in these supracellular assemblies. In agreement with this, overexpression of different Flag-tagged Zasp52 isoforms in stripes in the embryonic epidermis, using *enGal4*, led to junctional localisation of these isoforms (Suppl.Fig. S4 G-H’), confirming that Zasp52’s binding partners appear to be present in epidermal junctions throughout the embryo.

### Partial and complete loss of Zasp52 affects epithelial morphogenesis

Zasp52-GFP colocalised with several ubiquitously localised components of apical junctions within the embryonic epidermis, but its own localisation was limited to a subset of junctions and in particular to supracellularly coordinated junctions as shown above. In order to uncover what the molecular role of Zasp52 in these junctions was, we analysed the embryonic phenotypes, firstly in the zygotic mutant *zasp52^Δ^*, in more detail.

*zasp52^Δ^* embryos, in addition to the reduction in F-actin accumulation at the salivary gland placode boundary shown above (Fig. 2A-C), showed a slight disorganisation of the placode in that some groups of cells appeared more constricted (Fig. 4 A’, B’, magenta arrows) than in the control (Fig. 4C-C’’), and invaginated glands often showed an aberrant lumen (Fig. 4B’’ inset, compared to control in 4C’ inset, 4D, D’), indicative of defects in coordination of cell behaviours such as apical constriction during invagination. At mid to late stages of embryogenesis, *zasp52^Δ^* embryos showed occasional holes within the epidermis where the salivary glands previously invaginated into the embryo (Fig. 4 E compared to wild-type in F, magenta arrow).

**Figure 4.**
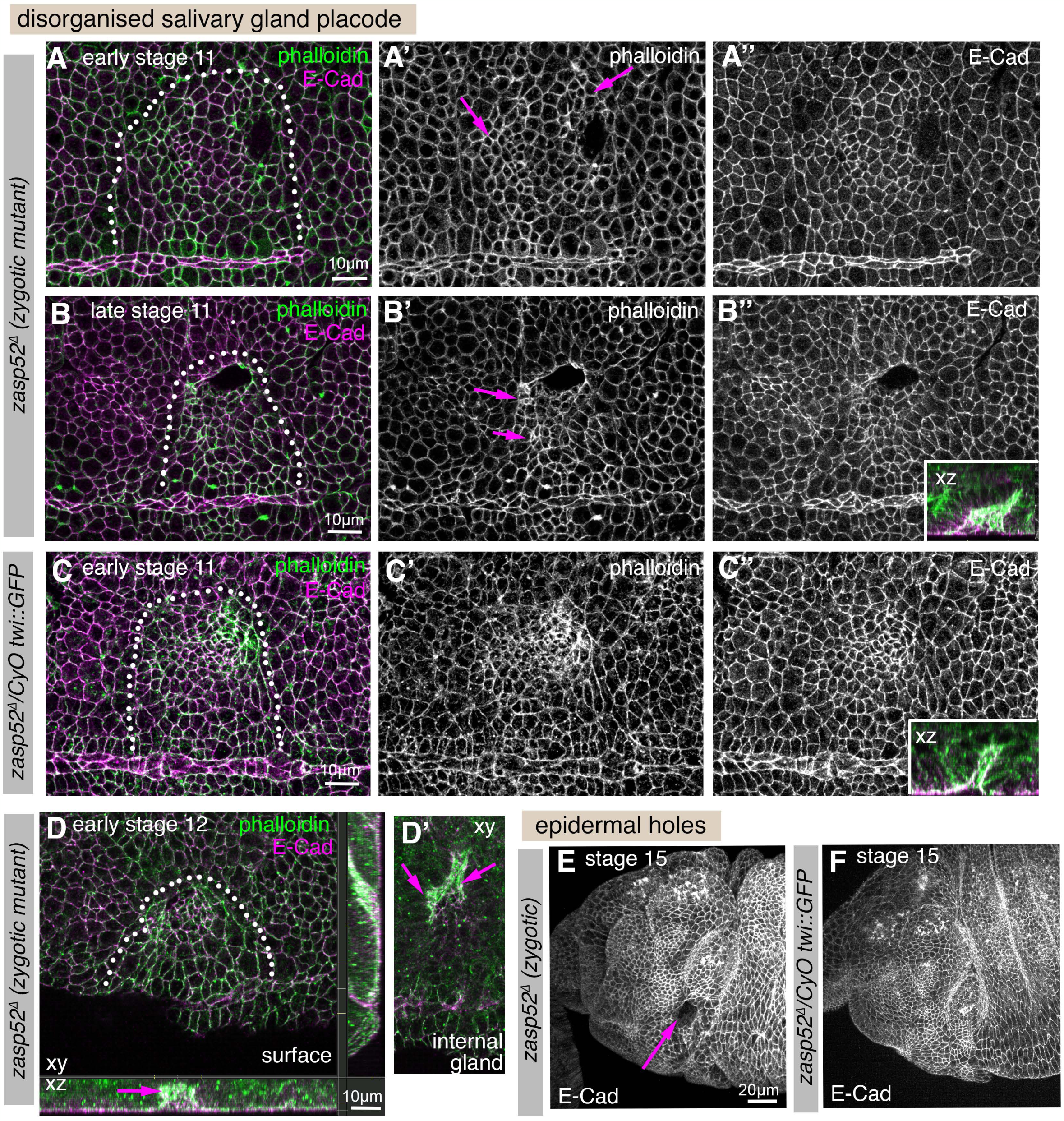
Zygotic loss of Zasp52 function leads to mild embryonic defects. **A-D** Salivary gland placodes in *zasp52^Δ^*(zygotic) mutant embryos at late stage 11 show slight disorganisation with the pattern of apical constriction (magenta arrows in **A’** and **B’**) not matching control placodes (**C-C’’**) and invaginated glands show aberrant lumen shapes already at stage 12 (**D**). E-Cadherin to label apical cell outlines is in magenta, phalloidin to label F-actin is in green. White dotted lines mark the boundary of the placode, xz and yz cross-section views (inset in **B’’, C’’** and in **D**) are indicated. **D’** shows the internal confocal sections below the epidermis of the placode shown in **D**. **E**, **F** At stage 15, when the salivary glands have completely invaginated, a third of *zasp52^Δ^* mutant embryos show an epidermal hole where the salivary gland would have previously invaginated from the surface (**E**). This is never seen in the wild-type (**F**). Labelling is E-Cadherin to mark apical cell outlines. Scale bars in **A-D’** are 10µm, in **E**, **F** 20µm.

As *zasp52* mRNA is provided maternally (https://insitu.fruitfly.org/cgi-bin/ex/insitu.pl; (Tomancak et al., 2002)), in order to observe the most severe possible phenotypes, we generated embryos lacking both maternal and zygotic Zasp52 contribution, *zasp52^Δm-/z-^* embryos. These embryos completely lacking Zasp52 displayed numerous defects, including a strong loss of patterned cell shapes in the early salivary gland placode compared to control placodes (Fig. 5 A-A’’ versus C-C’’) or even holes or tissue disruption where the placode is located (Fig. 5 B-B’’, arrows in B’, compared to D-D’’), as well as aberrant lumens of glands that managed to invaginate (Fig. 5B’’ inset versus D’’ inset; E, E’ at stage 14 compared to control in J, J’). Later stage embryos often showed wider epidermal disruption, in particular in regions where supracellular actomyosin cables were present in wild-type embryos. This was visible as the disruption or lack of ventral midline structures (already visible at stage 10 in Fig. 5A, arrows, and loss of the ventral midline at stage 15 in Fig. 5F). Furthermore, head involution, a process whereby the anterior-most structures of the embryo are internalised in major morphogenetic movements, appeared to partially fail, with structures that should have been internalised remaining on the outside along with significant epidermal tears visible in these regions (Fig. 5 F-I, white arrows pointing to non-internalised structures, magenta arrows pointing to epidermal holes and tears, compare to control in K). These phenotypes are reminiscent of disruption seen in weak alleles of *shg* (E-Cadherin) or alleles of alpha-Catenin that cannot interact with E-Cadherin and thus disrupt the link between E-Cadherin and actin (Orsulic and Peifer, 1996; Tepass et al., 1996), in line with Zasp52 also playing a role in the linkage of actin to junctions.

**Figure 5.**
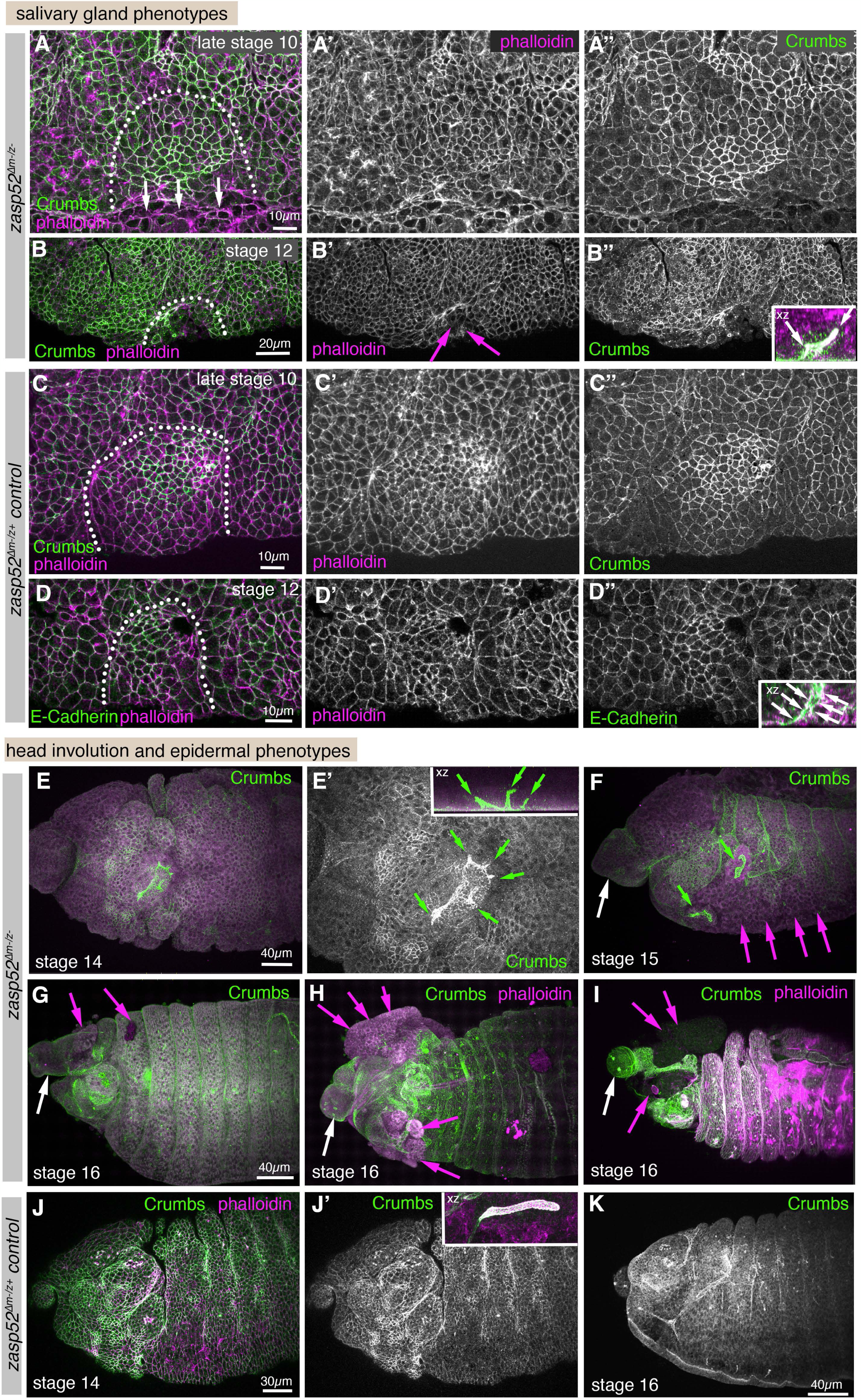
Complete loss of Zasp52 function severely disrupts morphogenetic processes. **A-D’’** Salivary gland placodes in *zasp52^Δm-/z-^* (maternal and zygotic) mutant embryos show disorganised salivary gland placodes with a disrupted boundary (**A**-**A’’**), and also epidermal tears in close proximity to where the supracellular actomyosin cable should be localised or where remnants of it remain (**B**-**B’’**, arrows point to tears). Wild-type placodes, by contrast show the graded pattern of apical constriction expected at late stage 10 as well as a smooth boundary (**C-D’’**). Invaginated salivary glands show highly aberrant lumens (**B’’**, inset, and **E’**, **F**), compared to the smooth narrow lumen of the control (**D’’** inset). White arrows in inset in **B’’** point to branched lumen, and to the narrow lumen in the control in **D’’**. Dotted lines mark the boundary of the placode. **E**-**K** *zasp52^Δm-/z-^* mutant embryos at later embryonic stages, stages 14-16, show severe problems with head involution, with major anterior parts failing to internalise (white arrows), compared to the control (**J**, **J’**) and furthermore tears and holes appearing in the epidermis though which internal structures protrude (magenta arrows) that are also absent in the control (**J**, **J’**). Also visible are the aberrant and branched lumens of the salivary glands (green arrows) compared to the narrow unbranched lumen in the control (inset in **J’**). Epidermal tears and holes were equally frequent in maternal zygotic as well as zygotically mutant embryos, and were paternally rescued (m^+^/z^-^: 4/12 embryos; m^-^/z^+^: 4/20 embryos; m^-^/z^-^: 12/35 embryos). Deformations due to failed head involution or loss of midline structures was very prevalent in maternal zygotic mutant embryos (24/35 m^-^/z^-^embryos) and only paternally rescued in half of these (7/20 m^-^/z^+^ embryos still showed the phenotype). Crumbs staining to label apical cell outlines is shown in green, and phalloidin to label F-actin in magenta. Scale bars are 10µm for **A**-**A’’, C-D’’**, 20µm for **B**-**B’’**, 30µm for **J**, **J’** and 40µm for **E-I** and **K**.

Therefore, loss of Zasp52 appeared to impair morphogenesis of tissues that displayed prominent supracellular actomyosin cables during their morphogenesis and led to large-scale disorganisation of embryonic tissues. The observed phenotypes of major aberrant tissue deformations as well as epidermal ruptures suggest that these cables could have served to coordinate major movements or protect tissue integrity.

### Genetic interaction suggests cooperation between Zasp52 and APC2

We identified above that Zasp52 is able to physically associate with several junctional components. We therefore generated a double mutant fly line lacking both Zasp52 and APC2, *zasp52^Δ^; apc2^D40^*, to analyse how this affected embryonic development.

Loss of APC2 alone within the embryonic epidermis, in *apc2^D40^* and *apc2^N175K^* mutant embryos, lead to phenotypes at the salivary gland placode boundary and at the leading edge-amnioserosa interface during dorsal closure that suggested an imbalance in forces of contracting cells with their neighbours (Suppl.Fig. 5). For instance, cells within the salivary gland placode appeared overconstricted, whereas in the epidermis just outside the placode boundary, cells were overstretched (Suppl.Fig. 5A-B’’). This could suggest a weakened adherens junction-actomyosin link in these mutants.

The embryos that are double zygotic mutant for Zasp52 and APC2, *zasp52^Δ^; apc2^D40^*, similar to the embryos lacking both maternal and zygotic Zasp52, displayed disrupted salivary gland placodal organisation (Fig. 6 A-B’, compare to C, C’), with cells displaying aberrant apical areas (green arrow in Fig. 6 A’), and embryos showing a disorganised ventral midline (Fig. 6 A’, B’, magenta arrows), and the dorsally located fold between maximillar and mandibular segments protruding erroneously into the placodal area (Fig. 6 A’, B’, white arrows). Furthermore, similar to the *zasp52^Δm-/z-^* mutant embryos, tears and holes appeared mid-embryogenesis around the ventral midline (Fig. 6 D, D’), and late in embryogenesis in the anterior part of embryos, again with head involution being impaired (Fig. 6 F, F’), compared to control embryos of matching stages (Fig. 6 E, G). In addition, the formation and closure of the amnioserosa and its surrounding actomyosin cable was affected (Fig.6 H-J’). At early stages during germband retraction, in some embryos, no clear boundary between the epidermis and amnioserosa developed (Fig. 6H’, magenta arrows, compare to 6I). In others, despite accumulation of F-actin at the leading edge front in many, though not all, cells the leading edge front was not taut, possibly due to a lack imbalance of tension (Fig. 6 J, J’, magenta arrows; compare to 6K).

**Figure 6.**
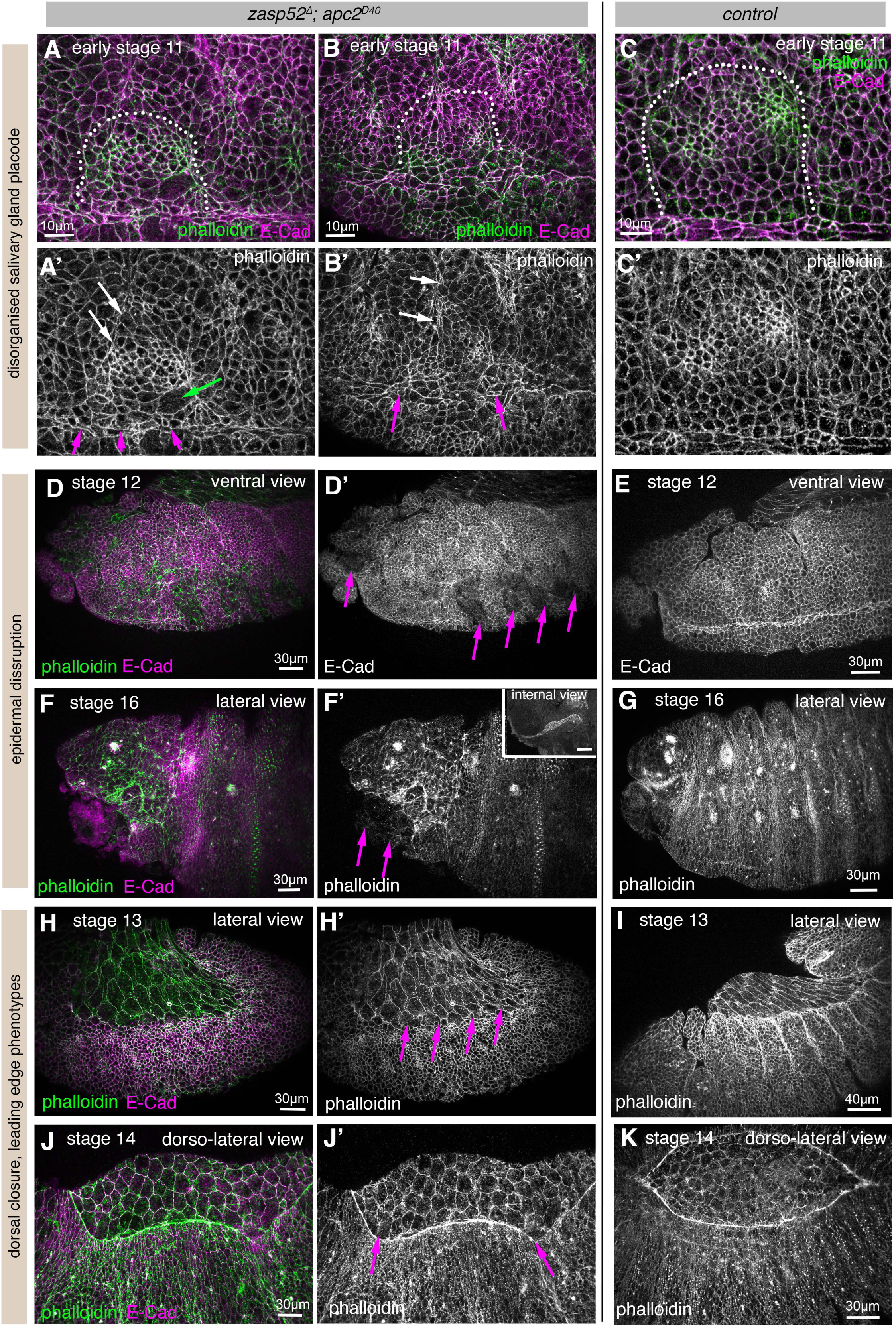
Zasp52 and APC2 interact genetically to strengthen supracellular actomyosin assemblies. **A-B’’** Salivary gland placodes in *zasp52^Δ^; apc2^D40^* mutant embryos show disorganised salivary gland placodes with aberrant patterns of apical constriction (green arrow), a disorganised boundary and disorganised ventral midline (magenta arrows) and the maximillar-mandibular fold invading the placodal area (white arrows), in comparison to control placodes (**C**, **C’**). Dotted lines mark the position of the boundary of the placode. **D-G** In *zasp52^Δ^; apc2^D40^* mutant embryos at stage 12 (mid-embryogenesis; **D**, **D’**) the epidermis near the ventral midline is often disrupted by tears (magenta arrows in **D’**), compared to the smooth intact epidermis in control embryos (**E**). At late embryogenesis, stage 16, the anterior head region in *zasp52^Δ^; apc2^D40^* mutant embryos shows tears and protruding internal tissues (**F**, **F’**; magenta arrows point to tears in **F’**), compared to the intact epidermis in control embryos (**G**). The invaginated salivary gland tube of the example embryo shown in **F**, **F’** also has an aberrant lumen shape (inset in **F’**). **H-I** Before dorsal closure commencing, *zasp52^Δ^; apc2^D40^*mutant embryos at early stages can fail to develop a clear boundary between epidermis and amnioserosa (magenta arrows in **H’**), compared to control embryos (**I**). **J-K** During dorsal closure *zasp52^Δ^; apc2^D40^* mutant embryos (**J, J’**) show uneven accumulation of F-actin in the leading edge cable as well as deformation of the cable indicative of loss of or uneven tension (magenta arrows in **J’**), compared to the homogeneous accumulation in control embryos (**K**). E-Cadherin labeling to mark apical cell outlines is in magenta, phalloidin to label F-actin is in green. Scale bars in **A-C’** are 10µm, in **D-H’, J, J’,** and **K** are 30µm, and in **I** is 40µm. See also Supplemental Figure S5.

In contrast to the milder phenotypes observed in *zasp52^Δ^*zygotic mutant embryos, the embryos double-mutant for *zasp52^Δ^*and *apc2^D40^* showed phenotypes more similar to those observed in *zasp52^Δm-/z-^* maternal and zygotic mutant embryos. This enhancement of the *zasp52^Δ^* zygotic mutant phenotype confirms what the co-immunoprecipitation results suggested, i.e. that Zasp52 and APC2 are working together to support junctions and junctional cytoskeleton involved in supracellular assemblies.

### Supracellular actomyosin cables containing Zasp52 act as mechanical insulators

Zasp52 as a component that is specifically enriched in embryonic supracellular actomyosin assemblies, in cooperation with other more widespread junctional components, appears to be important for the persistence of these cables and the morphogenetic events associated with them. It is still difficult, however, to pinpoint the collective function of such supracellular actomyosin cables during embryogenesis. Some functions for certain cables have been proposed, for instance for parasegmental cables in maintaining compartment identity against challenges such as cell divisions (Monier et al., 2010), or for the cable during dorsal closure in ensuring a taut and coordinated advancing front of epithelial cells (Ducuing et al., 2015). However, the function of the cable around the salivary gland placode, the cables at the ventral midline as well as the network of ventral anterior cables present during head involution have not yet been elucidated.

We suspect that one function of supracellular actomyosin cables is to insulate certain morphogenetic processes from others occurring nearby, in order to prevent undue physical interference between different morphogenetic events. Such effect could in fact be observed when we analysed the movement of cell nodes or vertices in the early salivary gland placode, when apical constriction at the position of the future pit is only just beginning at late stage 10 (Fig. 7 A-D). At this stage the actomyosin cable near the forming pit at the dorsal-posterior boundary of the placode is already visible (Fig. 7A-B’). Cell vertices that are inside the placode and near, but not part of, the forming invagination pit move towards the pit by about two micrometres over 10 min of apical constriction that has just commenced at the future pit position (Fig. 7 A’’, B’’, C and D; vertices of actively constricting cells marked in green in A and B were excluded from the analysis as these moved due to the apical constriction). By contrast, cell vertices equally close to the forming pit but posterior to the actomyosin cable surrounding the placode, and hence outside the placode, barely move at all (Fig. 7 A’’, B’’, C and D). These observations indicate that one function of the actomyosin cable surrounding the salivary gland placode might be to serve as a barrier to insulate morphogenetic movements within the placode from the surrounding epidermis.

**Figure 7.**
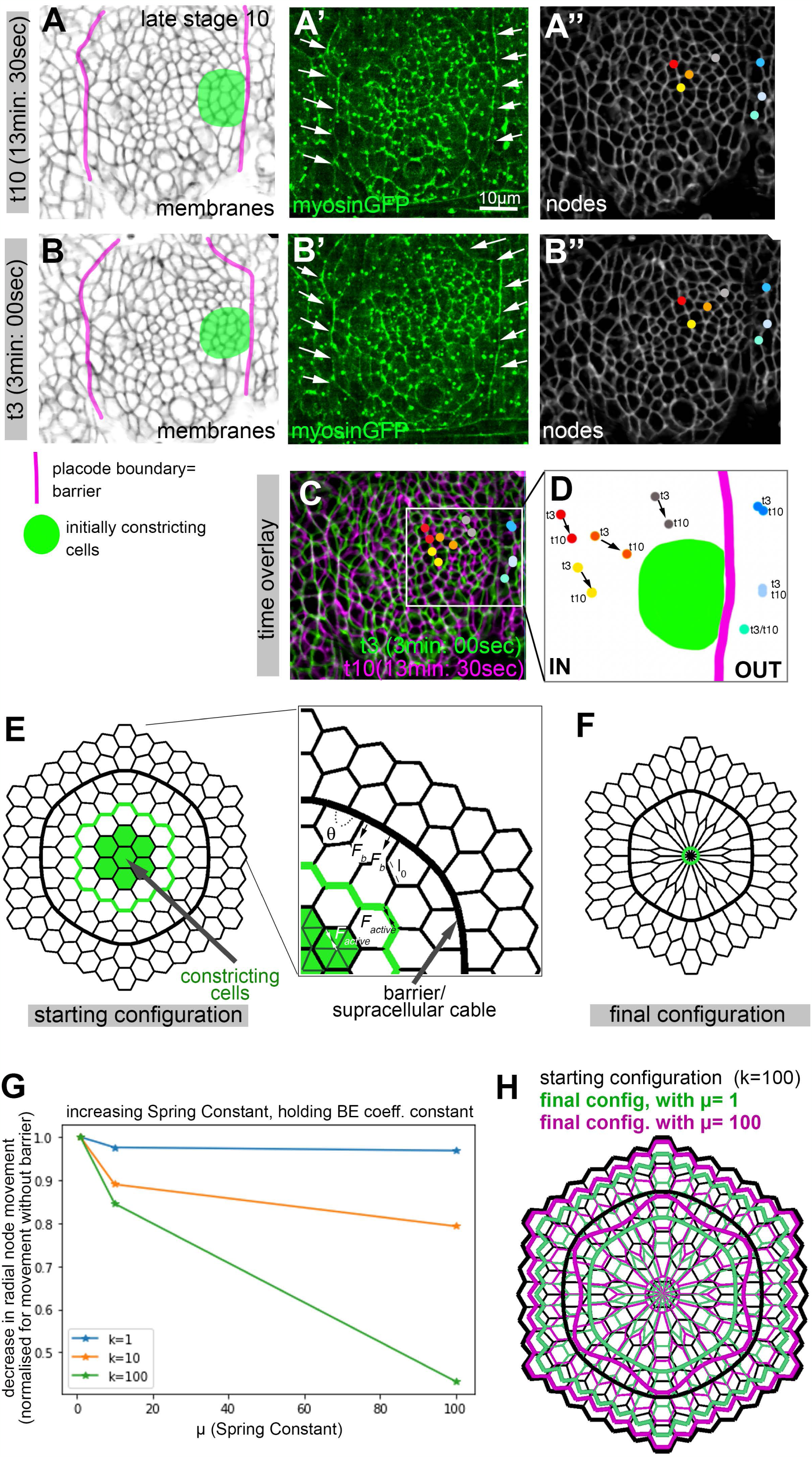
In vivo analysis and in silico modeling of the effect of a rigidity and contractility barrier on cell vertex movement. **A-D** Qualitative analysis of cell vertex movement of vertices near the forming invagination pit where apices constrict (as in the model), for vertices inside or outside the actomyosin cable. **A, B** Stills from a time lapse movie, 10min:30sec apart (t3 and t10 of the movie), with the placodal boundary where the actomyosin cable is positioned marked in magenta and the initial group of constricting cells at late stage 10 marked by green. **A’, B’** Still of the same movie showing the SqhGFP channel to show myosin localisation. Arrows point to the position of the cable. **A’’, B’’** Nodes near the constriction zone inside and outside the actomyosin cable are marked by coloured dots at t3 (**A’’**) and t10 (**B’’**). **C** shows both time points false coloured and superimposed (t3 in green and t10 in magenta, with the highlighted vertices shown for both timepoints). **D** Close up of the positions of the vertices inside and outside the cable an t3 and t10. The position of the cable is marked in magenta and of the constricting cells in green. **E-H** A vertex model of a simplified symmetrical 2D version of the placode was implemented similar to (Durney and Feng, 2021). **E** The starting configuration assumes individual cells as hexagons apart from cells next to a barrier or supracellular cable (thick black line) where due to an implemented contractility and hence rigidity, as well as a bending energy, cell junctions of the barrier are aligned (as is seen *in vivo*) with tricellular junctions showing 90°, 90°, 180° angles. A central group of 7 cells as well as some selected edges (green) constrict to represent the forming invagination pit, and movement of nodes outside the barrier (representing the epidermis surrounding the salivary gland placode) is assessed. **F** shows an example of a final configuration. **G** Decrease in radial movement observed for the outside nodes, normalised by the movement without the barrier, for different values of a bending energy coefficient k (that reflects a penalty for the nodal angles deviating from 120° at the boundary) observed for increasing spring constants µ. **H** shows a comparison of the starting configuration compared to two different final configurations, with k=100 for both and µ= 1 (green configuration) or µ=100 (magenta configuration). See Supplemental Movie S1.

In order to explore whether ‘morphogenetic insulation’ could be a general role of actomyosin cables containing Zasp52 and what properties could confer such functionality, we employed an *in silico* 2D vertex model of the salivary gland placode to test this (Fig. 7; (Durney and Feng, 2021)). This model considers a centre of hexagonal cells surrounded by a barrier of increased contractility and stiffness to model the actomyosin cable surrounding the placodal cells, with a further two coronae of cells surrounding the barrier that represent the surrounding epidermis (Fig. 7E). In this model, the central cells constrict, thereby pulling on and deforming other cells on the inside and outside of the barrier (Fig. 7F). The starting state takes into account that junctions at the barrier *in vivo* are more aligned and hence vertices at tricellular junctions deviate drastically from 120°. This is reflected in the model by the contribution of a bending energy coefficient (k) on edges representing the supracellular barrier. The central cells are allowed to actively constrict and the average nodal movement towards the centre is measured, whilst the spring constant (µ) of the barrier is varied. For a set value of the bending energy coefficient k (1, 10 or 100) an increase in spring constant µ consistently led to a decrease in the nodal movement of cells outside the barrier towards the centre (Fig. 7G and H).

Thus, the *in vivo* and *in silico* data above suggest that during mid to late embryogenesis in the fly the many large-scale actomyosin cables observed act as insulators, thus ensuring that morphogenetic movements do not spread beyond a defined primordium, or act to prevent neighbouring regions undergoing different morphogenetic processes from unduly influencing each other.

## Discussion

The formation of the physical shape and functionality of organs during development requires coordination at many different time and length scales. In many cases cell behaviours between groups of neighbouring cells within a tissue primordium need to be coordinated. Conversely, at boundaries between differently fated embryonic regions or organ primordia, such coordination has to cease so as not to inadvertently affect the neighbouring processes. Coordination between groups of cells is often achieved through coordination of the cells’ cytoskeletal systems, be it alignment of actin or of microtubules (Röper, 2013; Sanchez-Corrales and Roper, 2018). Any coordination of these systems is transmitted and coupled via cell-cell adhesion complexes usually located at adherens junctions.

One very obvious coordination of the cytoskeleton observed during numerous morphogenetic events in both invertebrates and vertebrates is found in the seemingly supracellular arrangement of actomyosin into supracellular cables. In *Drosophila*, numerous actomyosin cables are formed throughout embryogenesis, from early parasegmental cables and cables flanking the midline to cables surrounding tissue primordia at later stages. During the morphogenesis of regions of the head including the salivary glands, an intricately cross-linked system of actomyosin cables develops (Röper, 2013). Though some studies have offered glimpses of functions or mechanisms of assembly for particular cables (Hashimoto and Munro, 2019; Major and Irvine, 2006; Monier *et al*., 2010; Pare et al., 2019; Pare et al., 2014; Röper, 2012; Sidor *et al*., 2020), major questions about their assembly, composition and general function remain. Though seemingly supracellular, individual cable segments in a cell are of course connected to neighbouring parts at cell-cell junctions. But are these junctions in any way different to regular adherens junctions? Furthermore, are there specific components of such supracellular cables that set them apart from ubiquitous cortical actomyosin?

Zasp52 as a core component of Z-lines in muscles is the first component of supracellular actomyosin cables that appears to set them apart from regular junctional actomyosin. We identified an actin-binding motif in Zasp52’s central domain, as well as a host of junctional proteins that interact with Zasp52. Moreover, we found that its N-terminus, known for binding actin (Liao *et al*., 2020), also promotes Zasp52 dimerisation and multimerisation. These results suggest that upon binding to Zasp52, junctional actomyosin likely takes on a new and high-order structure. It is noteworthy that the junctional interaction partners localise to the marginal zone (Crumbs, Patj), the adherens junctions (armadillo, alpha-Catenin, p120-Catenin, Shotgun/E-Cadherin, APC2, Sidekick, Bazooka, Pyd/ZO-1, Canoe/Afadin) and even basally to adherens junctions (Scribble). This suggests that the increased size of actomyosin cables, which can be inferred visually from the strongly increased intensity of F-actin and myosin labelling in these structures, means that they span an enlarged region along the lateral membrane compared to general junctional actomyosin that is usually restricted to adherens junctions only (Desai et al., 2013; Röper, 2015). Such lateral expansion of actomyosin accumulation could be a defining feature of supracellular actomyosin structures, with a possible actomyosin organisation across the epithelial junctions similar to lateral stacking of thin filaments in sarcomeres, though this requires future investigation.

The phenotypes observed in the absence of Zasp52, as well as in the combined absence of Zasp52 and APC2 (but not APC2 alone), in particular the epidermal tears and holes, are reminiscent of phenotypes observed when adherens junctions and their linkage to the actin cytoskeleton are compromised by the lack of alpha-Catenin (Sarpal et al., 2012), or by hypomorphic mutations in E-Cadherin itself (encoded by the *shotgun* (*shg*) locus; (Tepass *et al*., 1996)). Embryos lacking alpha-Catenin or certain hypomorphic classes of *shg* mutant embryos also show epidermal holes, especially in the head and ventral region, as well as anterior disorganisation that might be a result of head involution defects. Thus, impairment of the linkage of adherens junctions to the junctional actin cytoskeleton, occurring in alpha-Catenin, E-Cadherin and also Zasp52 mutants, appears to lead to a common set of phenotypes.

Interestingly, several of the junctional components identified in the complex with Zasp52 are either enriched at or are exclusive to tricellular junctions (APC2, Canoe/Afadin, Sidekick), and Zasp52 is enriched in these tricellular junctions itself. These cell vertices are the points where individual segments of actomyosin cables require to be connected to the next cable segment one cell on. The presence of Zasp52 in precisely these junctions could help to reinforce them against the increased tension that actomyosin cables usually exert and bear (Fernandez-Gonzalez et al., 2009; Röper, 2012).

Recruitment of Zasp52 to a forming cable might well occur through a positive feed-back loop. Initial Zasp52 recruitment to increased junctional F-actin at a position where a cable is forming, for instance at the boundary of the placode (downstream of Crumbs anisotropy and Rok accumulation; (Röper, 2012; Sidor *et al*., 2020)), could structurally change the cable. This could be via strengthening it through actin crosslinking, or via strengthening and amplifying the connection to junctions, which in turn could promote further actin and myosin recruitment, as well as increased recruitment of Zasp52 itself. Actomyosin enrichment, for instance in the salivary gland placodal cable, occurs before Zasp52 recruitment to the same location. Thus, it seems to be the localisation of actomyosin, not of Zasp52, that determines where a cable forms. Rather, actomyosin cables containing Zasp52 might assemble at the coincidence of two events: firstly, the enhanced accumulation of junctional actomyosin triggered by other mechanisms, and secondly, the expression of Zasp52 in the very same cells, thereby leading to its recruitment via interactions with junctional binding partners to the forming cable. This recruitment then adds additional cable characteristics. Zasp52’s role in altering the assembly or structure of a cable could be even more profound, as its role in Z-lines of sarcomeres could suggest that actomyosin cables might have a higher order of actin and myosin assembly than general actomyosin, imposed by recruitment of Zasp52. Such higher order could be observed previously in specialised junctions of support cells in the mouse cochlea, as a feature of fully differentiated cells rather than a tool of morphogenesis (Ebrahim et al., 2013). The loss of actin accumulation at cable sites in the *zasp52* mutant embryos strongly suggests that actin stabilisation could be a key part of Zasp52’s function. Dissecting these structural changes in detail will be the next challenge in understanding actomyosin cable function.

Can our analysis of the phenotypes observed in *zasp52* mutants furthermore allow us to identify a more general role for actomyosin cables during morphogenesis? One key role could be that the higher stiffness and contractility of actomyosin cables allows them to serve as physical barriers, thereby insulating morphogenetic events and preventing them to unduly influence surrounding tissues. Such undue spread of movement can easily be imagined as all epithelial cells will be mechanically coupled to neighbouring cells to some degree due to their epithelial junctions. Hence, defined mechanical barriers could be an important requirement during epithelial morphogenesis. To assess this option we turned to the properties of the actomyosin cable surrounding the salivary gland placode. Our combined *in vivo* and *in silico* approaches demonstrate that the placode cable is acting as a mechanical insulator, which could hence also be true for other cables. A further role could be that the presence of interlinked cables spanning large regions of the epidermis allows a coordination of large-scale movement of different tissue primordia. In the *Drosophila* embryo, this might well be the case for the interconnected cables in the ventral head region. And matching this, the *zasp52* complete null mutant shows major defects in this process. More detailed whole embryo studies in the future will reveal how common these function for actomyosin cables are.

In summary, the analysis of Zasp52 as a component of actomyosin cables has revealed what we suspect are key functions of actomyosin cables in epithelial morphogenesis in animals: the mechanical insulation of individual morphogenetic events as well as the coordination of large-scale epidermal changes. It will be interesting to analyse whether these functions of Alp/Enigma proteins are conserved in other prevalent cables observed during vertebrate morphogenesis.

## Material and Methods

### Key resources table

| REAGENT or RESOURCE | SOURCE | IDENTIFIER |
| --- | --- | --- |
| **Antibodies** |  |  |
| Anti-Apc2 (rabbit) | http://dx.doi.org/10.1038/11064 | N/A |
| Anti-Crumbs (mouse) | Developmental Studies Hybridoma Bank at the University of Iowa (DSHB) | DSHB Cat#Cq4; RRID: AB_528181 |
| Anti-E-Cadherin (rat) | Developmental Studies Hybridoma Bank at the University of Iowa (DSHB) | DSHB Cat#5D3; RRID: AB_528116 |
| Anti-FLAG (mouse) | Sigma | Cat#M2 |
| Anti-GFP (rabbit) | Abcam | Cat#ab290 |
| Anti-Patj (guinea pig) | http://dx.doi.org/10.1083/jcb.201206064 | N/A |
| Anti-Sidekick (guinea pig) | http://dx.doi.org/10.1242/dev.158246 | N/A |
| Anti-Zasp52-N (rabbit) | http://dx.doi.org/10.1083/jcb.200707045 | N/A |
| Donkey anti-Goat IgG (H+L) Cross-Adsorbed Secondary Antibody, Alexa Fluor™ 488 | Invitrogen™ | Cat#A-11055 |
| Goat anti-Rabbit IgG (H+L) Cross-Adsorbed Secondary Antibody, Alexa Fluor™ 488 | Invitrogen™ | Cat#A-11008 |
| Goat anti-Guinea Pig IgG (H+L) Highly Cross-Adsorbed Secondary Antibody, Alexa Fluor™ 647 | Invitrogen™ | Cat#A-21450 |
| Goat anti-Mouse IgG (H+L) Cross-Adsorbed Secondary Antibody, Alexa Fluor™ 488 | Invitrogen™ | Cat#A-11001 |
| Cy™5 AffiniPure Donkey Anti-Mouse IgG (H+L) | Jackson ImmunoResearch Laboratories | Cat#715-175-151 |
| Alexa Fluor® 647 AffiniPure Donkey Anti-Rat IgG (H+L) | Jackson ImmunoResearch Laboratories | Cat#712-605-153 |
| **Bacterial and virus strains** |  |  |
| Library Efficiency™ DH5α Competent Cells | Invitrogen™ | Cat#18263012 |
| BL21(DE3) Competent Cells | Thermo Scientific™ | Cat#EC0114 |
| **Chemicals, peptides, and recombinant proteins** |  |  |
| GST | Joseph Watson | N/A |
| Rabbit muscle actin | Cytoskeleton | Cat#AkI99 |
| Rhodamine-phalloidin | Cytoskeleton | Cat#PHDR1 |
| Vectashield | Vectorlabs | Cat#H-1000 |
| **Experimental models: Cell lines** |  |  |
| S2 cells | S2-DRSC (DGRC Stock 181 ; https://dgrc.bio.indiana.edu/stock/181) | RRID:CVCL_Z992 |
| **Experimental models: Organisms/strains** |  |  |
| *Drosophila melanogaster: apc2<sup>D40</sup>* | Drosophila Bloomington Stock Centre | Cat#6801 |
| *Drosophila melanogaster: apc2<sup>N175K</sup>* | Drosophila Bloomington Stock Centre | Cat#7210 |
| *Drosophila melanogaster: actnCC01961* | Drosophila Bloomington Stock Centre | Cat#51573 |
| *Drosophila melanogaster: enGal4* | Drosophila Bloomington Stock Centre | Cat#1973 |
| *Drosophila melanogaster: Df(2R)BSC308* | Drosophila Bloomington Stock Centre | Cat#23691 |
| *Drosophila melanogaster: hsFLP; ovoD1 /TTP* | Gift from Isabel Palacios | N/A |
| *Drosophila melanogaster*: shgTdTomato | Drosophila Bloomington Stock Centre | Cat#58789 |
| *Drosophila melanogaster*: sdk<sup>MB05054</sup> | Drosophila Bloomington Stock Centre | Cat#24603 |
| *Drosophila melanogaster*: sqh-TagRFPt[3B] | https://dx.doi.org/10.1016%2Fj.devcel.2019.05.027 | N/A |
| *Drosophila melanogaster*: zasp52<sup>Δ</sup> | Drosophila Bloomington Stock Centre | Cat#59018 |
| *Drosophila melanogaster*: zasp52-GFP[G00189] | Drosophila Bloomington Stock Centre | Cat#6838 |
| *Drosophila melanogaster*: zasp52-GFP[ZCL423] | Drosophila Bloomington Stock Centre | Cat#58790 |
| *Drosophila melanogaster*: armadillo-YFP[CPTI001198] | Kyoto Stock Centre (DGGR) | Cat#115-134 |
| *Drosophila melanogaster*: UAS-Zasp52-PK-6xHis-Flag | Kind gift of Frieder Schöck; (Liao *et al.*, 2020) | N/A |
| *Drosophila melanogaster*: UAS-Zasp52-PRΔPDZ-6xHis-Flag | Kind gift of Frieder Schöck; (Liao *et al.*, 2020) | N/A |
| **Oligonucleotides** |  |  |
| Zasp52-PF actin binding site (FseI and Ascl sites introduced) residues 1312 – 1319)<br>(see details below due to length) | Synthesised by Twist Bioscience | N/A |
| Zasp52-PF (FseI and Ascl sites introduced) residues 1 – 2198)<br>(see details below due to length) | Synthesised by Twist Bioscience | N/A |
| Zasp52-PF PDZ + ZM (FseI and Ascl sites introduced) residues 1 – 216)<br>(see details below due to length) | Synthesised by Integrated DNA Technologies | N/A |
| **Recombinant DNA** |  |  |
| pGEX dElp4S FA D338 | kind gift of Emmanuel Derivery;<br>https://doi.org/10.1038/nature16443 | N/A |
| pMT puro FA | kind gift of Emmanuel Derivery;<br>https://doi.org/10.1038/nature16443 | N/A |
| **Software and algorithms** |  |  |
| Clustal Omega | https://doi.org/10.1093/nar/gkac240 | N/A |
| ImageJ/Fiji | NIH | N/A |
| Imaris | Bitplane | N/A |
| Perseus (MaxQuant) | https://doi.org/10.1038/nmeth.3901 | N/A |
| Prism | Graphpad | N/A |
| Vertex Based Model of Epithelial Mechanics (in house) | DOI: 10.5281/zenodo.7016948<br>(this publication) | N/A |
| **Other** |  |  |

### Fly husbandry and genetics

*Drosophila* embryos were collected for staining on apple juice agar plates at 25°C for overnight or day durations as indicated. Embryos used for live imaging were collected at 25°C and aged 2 hrs at 29°C prior to imaging. Embryos for immunoprecipitations were pre-laid on 90 mm apple juice agar plates for 1 h to avoid collecting older retained embryos, the plate was discarded and collections started for 9, 10, 11 or 12 hrs. Overnight collections without pre-lay were collected in the same manner.

Genotypes used in figure panels:

**Table 1.**
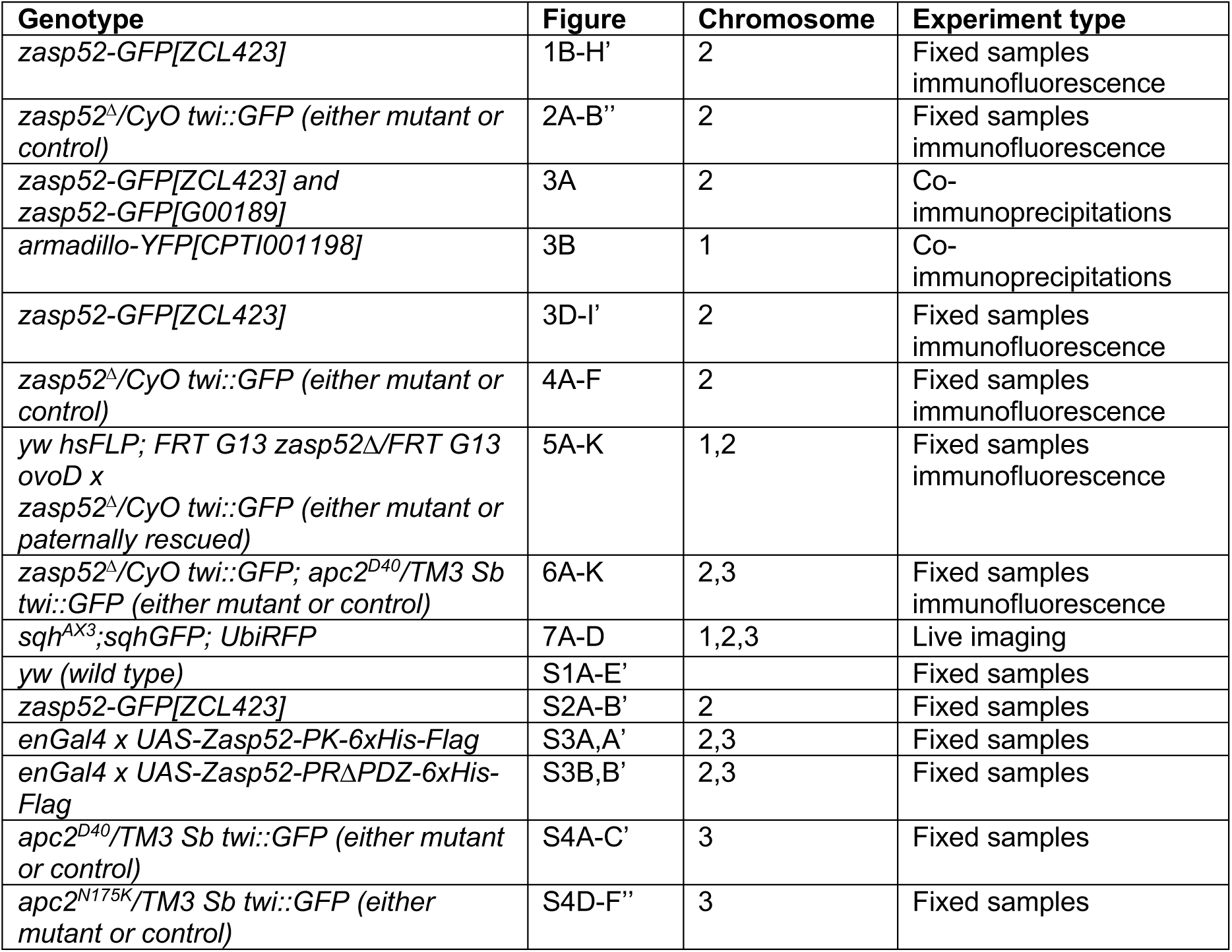
Embryo genotypes as presented in figures.

### Generation of zasp52^Δ^ germline clones

To generate embryos devoid of maternally deposited Zasp52, the FLP-DFS strategy was employed, based on the work of Chou and Perrimon (Chou and Perrimon, 1992). To this end, *FRT G13 zasp52^Δ^* virgins were crossed to males of *hsFLP; FRT G13 ovoD1 /TTP* stock. Progeny of this cross was heat shocked for 1 h at 37°C when reaching first instar stage and again the following day. From these crosses, virgin females were picked and crossed to *zasp52^Δ^/CyO twi::GFP* males and embryos collected and analysed.

### Immunofluorescence staining of whole-mount embryos

Embryos of desired stages were collected from apple juice-agar plates with a brush and tap water and transferred into a sieve. After rinsing with tap water, embryos were dechorionated with 50% of thick bleach/water for 3 min. Embryos were washed several times with water to remove bleach solution and residual water absorbed with paper wipes. Embryos were then taken up in 800 µl heptane and placed in a small glass screw-cap vial. 400 µl PBS and 400 µl 8% PFA (EM-grade)/PBS were added to the embryos and embryos left to fix on a shaker for 15 min. The lower phase containing PFA was removed and embryos devitellinised in 1 ml 90% EtOH/H2O with shaking. The embryos were stored at –20°C in EtOH.

Devitellinised and fixed embryos were simultaneously permeabilised and blocked with PBT (0.3% Triton X-100, 2.5% BSA in PBS) for one to three hours at 4°C. Primary antibodies were diluted in PBT and embryos incubated with antibodies overnight at 4°C on a shaker. Embryos were washed three times with 500 µl PBT for 20 min on shaker at RT. Secondary antibodies were diluted 1:200 in PBT or according to manufacturer and added to washed embryos to incubate 2.5 hours at RT or overnight at 4°C. Embryos were subsequently washed 3x with PBT for 20 min on shaker at RT. Embryos were mounted in Vectashield mounting medium (H-1000).

### Live Imaging of whole-mount embryos

Embryos were collected from apple agar plates, dechorionated and maintained in water until mounting. Embryos were oriented on agar to the desired position and attached to a cover slip prepared with heptane glue. The cover slip was pushed onto sticky metal slides and embryos covered with halocarbon oil 27. The embryos were then kept at 25 or 29°C depending on the stage to be imaged. Image collection was performed on a Zeiss 780 UV confocal microscope and spectral unmixing applied when imaging GFP or YFP due to autofluorescence of the embryo.

### Ectopic expression of Zasp52 in engrailed stripes

Zasp52 truncations were expressed with the UAS-GAL4 system using the *enGal4* driver. To this end, virgins of *+/+; enGAL4/CyO, UAS-GFP* were crossed with *w*/Y; UAS-Zasp52-PK-6xHis-FLAG/TM3 twi::GFP* or *w*/Y; +/+; UAS-Zasp52-PRΔPDZ-FLAG/TM3 twi::GFP*, and the resulting F1 embryos fixed and stained with anti-Cadherin and anti-FLAG antibodies as well as with rhodamine-phalloidin. Fluorescent images of engrailed stripes in the ectoderm were acquired with on an Olympus FluoView 1200.

### Quantification of F-actin signal intensity at the supracellular actomyosin cable in the placode

Whole-mount embryos were fixed and stained as described above. Higher resolution images of placodes were acquired on the Olympus FluoView 1200 microscope as z-stacks. Equivalent slices (8 slices corresponding to 400 nm, starting apically at first observed intensity of E-Cadherin) were used to create a SUM projection. Three rectangular areas were defined over the whole placode to measure background intensity and calculate average background intensity. Phalloidin signal intensity of individual junctions at the placode border were measured, leaving out the two rows of cells at the ventral midline and the junctions closest to the invagination pit because of distortion of the tissue here. Junction intensity was normalised to background ((IJunction/AreaJunction)/(Ibkgd/Areabkgd)).

### Transfection and staining of S2 cells with Zasp52 truncations

S2 cells (UCSF, mycoplasm-free judged by DAPI staining) were grown at 25°C in Schneider’s Medium (ThermoFisher) supplemented with 1% Pen/Strep (Gibco) and 10 % (vol/vol) Fetal Bovine Serum (Gibco; Heat-inactivated for 1h at 70°C). Stable cell lines were obtained by transfection with pMT puro vectors containing either Zasp52-PF or Zasp52-PF-ABM using Effectene (Qiagen) according to the manufacturer’s instructions, followed by selection in 5 µg/ml Puromycin (ThermoFisher). Expression was induced with 0.6 mM CuSO4 (in selection medium) for two days. Ibidi chambers were coated with 50 µg/ml Concanavalin A (diluted in PBS) for 10 minutes and washed with PBS. Cells were added in medium to the wells and allowed to spread for 1h. Cells were washed with PBS, fixed with 4% PFA (diluted in PBS) for 20 minutes, permeabilised with Triton-X100 (0.1% in PBS), washed and stained with primary antibody (diluted in 0.1% BSA-PBS) for 1h. Cells were washed with PBS and stained with secondary antibody (diluted in 0.1% BSA-PBS) for 1h, washed and imaged directly in PBS. Images were acquired on a Zeiss 880 confocal microscope.

### Recombinant expression and purification of the central Zasp52 actin binding motif

The Zasp52 actin binding motif was expressed as a GST fusion protein in *E. coli* and purified via affinity and size exclusion chromatography. The plasmid was generated via synthetic assembly of aa1131 – aa1318 of the Zasp52-PF cDNA (synthetic assembly of Zasp52-PF cDNA, Twist Bioscience) with AscI and FseI restriction enzyme overhangs and was digested and ligated in pGEX F/A vector cut with AscI and FseI. The pGEX plasmids containing truncations were transformed into the Rosetta (DE3) strain via heat shock transformation and selected via antibiotic selection. Colonies were picked the next day to inoculate pre-cultures. Precultures were used to set-up large cultures grown to OD600 = 0.6 and expression was induced by addition of 0.5 mM IPTG. Cultures were grown overnight at 18°C and harvested the next day via centrifugation at 4000Kxg for 30 min. Supernatant was discarded and pellets lysed on ice by resuspending in lysis buffer (50 mM Tris-HCl pH 7.4, 150 mM KCl, 1% Triton X-100, 10 mM MgCl2, 5 % glycerol, 1 mM DTT, 1x complete protease inhibitors, 0.7 mg/ml lysozyme and 10.00 µg/ml DNAse I in water) and then sonicated on ice followed by clearing of lysate via centrifugation at 60,000xg for 30 min at 8°C. Protein-containing supernatant was incubated with 1 to 5 ml of GSH-beads equilibrated in lysis buffer on a shaker for 2 hours at 4 °C. Flow-through was collected and the column was washed once with 45 CV of wash buffer (50 mM Tris-HCl pH 7.4, 150 mM KCl, 5 % glycerol, 1 mM DTT). To remove chaperones, an additional wash step with 1 CV ATP wash buffer (50 mM Tris-HCl pH 7.4, 150 mM KCl, 5 % glycerol, 10 mM NaATP, 10 mM MgCl2, 1 mM DTT) was carried out. GST-fusions were eluted with 10 mM glutathione (100 mM Tris-HCl pH 7.4, 150 mM KCl, 10 mM GSH, 5 % glycerol, 1 mM DTT) in 1 CV fractions. Fractions rich in GST-fusion protein were pooled and cleaved using TEV enzyme during dialysis (50 mM Tris-HCl pH 7.4, 150 mM KCl, 5 % glycerol, 1 mM DTT) overnight at 4°C. In order to remove free GST from the solution, the dialysed protein was incubated with equilibrated GSH resin for 2 hrs at 4°C and the flow through collected, concentrated and injected in a Superdex 200 (16/100, GE Healthcare) exclusion chromatography column. The column was equilibrated with gel filtration buffer (20 mM K-HEPES pH 7.5, 150 mM KCl, 5% glycerol, 1 mM DTT). Fractions collected were analysed by staining on SDS PAGE and fractions chosen, pooled and concentrated. Aliquots were frozen in liquid nitrogen and stored at -80°C until used for experiments.

Zasp52-PF actin binding site (FseI and AscI sites introduduced) residues 1312 – 1319): ggtgggccggccagagcctcgattgtgtctgccttgaaggaggaaaccgatctggagtaccagaagtatctcaaggcccagcagcgcaaccagaaaagattggactacttccaccagaaagaggaggagctctcgggtctgcagggccaacagctaacccaacttcagagggagctctcgaaccagcaacagaatcttctgagccaacagcaactgcagcaatccaagttgttgcaactgcagcagtgcgtccagagccaagagttgcagcaacaggtgcagcatctcacccagaaatcacaacagcaacctcctcaagctaaccaacagcagcaacaacagcaacagcaacggggtacccaacagcagcaacactcccaagtaacccaaagaacccaacagcaacagcaacaagtgccccaacaagtaacccaacagcaacaacaagaacactctctgctatcgcaaaccacactcgctgagacccaaacccttcaggccaatgcccagtctcagagttccgcttcctacagctccaaagcgactgcttgctctaactcctcttccacagtcggcgcgccggtg

Zasp52-PF (FseI and AscI sites introduduced) residues 1 – 2198): atgcggccggcccatggcccaaccacagctgctgcaaatcaaattgtcacgtttcgatgcccaaccctggggattccgccttcaggggggcacggacttcgctcagcccctgctggtgcaaaaggtgaacgccggcagcttgtccgagcaggctggcctccagcccggcgatgcggtggtcaagatcaatgacgtggatgtcttcaatctgcgtcacaaggatgcccaggacattgtggtgcgctccggcaacaactttgtcatcacagtgcagcgcggtggctccacctggcgcccgcatgtgacaccgactggcaatgtgccgcagcccaactcgccgtatctgcagacggtgacgaagacctctct ggctcacaaacaacaggacagccagcacatcggctgtggctacaacaacgcggcccgtcccttctccaacggcggcgatggcggcgtgaagagcattgtcaataaacaatacaacaccccggttggcatttacagcgatgaatctattgcggaaacactctcggcccaggcggaggttttggctggcggtgtgctcggcgtcaacttcaagaagaacgagaaggaataccagggcgatcgctccgaggttctgaagttcctgcgcgaggaggagaccggccagtccactccagcattcggcaatagccactacgagcatgatgcaccacagcaactgcaacagccacaacagcaatacaaccaacaccagcaacactatcaccagcaacaacaacaacagcaatcgagcaccactcgccatgtcagcgcccccgtgaactcccccaagcccccgagcaccggcggactcccaactggccagaacatttgcaccgaatgcgagcgcctcattactggcgttttcgtgcgcatcaaggataagaacctgcacgtggagtgcttcaagtgtgccacgtgtggcacctcgctgaagaaccagggctactacaacttcaacaacaagctctactgcgacatccacgccaaacaggccgccatcaacaatccccccaccggcaccgagggctacgtccccgttcccatcaagcccaacaccaagctgagtgcctccaccatctcatcggccttgaactcgcacggatacggtggccactcgaacggctactccaatggaaactccacccctgctccggcaccggttgcaagctctcaagcaacagcaacagtagcaacggtagcaccatccgctgcaacagcagcaactgcagcagcaacaccccaagcagcaactgcaacagatagcccagctgcaacagcatcatcatcagacaatatgtcggcctacgtggcagatgagccctcttcgatttatggccaaattagcgctgaatcggtggcattggccccaccaccaccacagccacccactgccggcgggggcgatcagccctttgagtacgtcacgctcaccggcaacgtcatccgcagcgtgcaggctcccggaaagggggcgtgccccagctacaaggtgaaccagggctatgctcgtccgttcggtgccgccgctcccaagtcgccggtgtcgtatccgccgcagcagcaacagcagtcgccgcgtcccgctcccggtggccaaaacccgtacgccaccctgccccgcagcaatgtgggccaacaaggtggagaggctgtggaggaactgcagccggaattcgaggaggaggattgctatgagatggacatcgaggtggccctggccgcaagtcgccaatcgcagcgtggctctagtttcacttggccaccgccgcaggatgatagccacttggcacccaccgcggcgcctctttatatccctccaccggagacgcaacatgtggtggtttcgaatccggtgcagcaagtgcctccattgccacctggaggagcaactgctcgactagatccgcaacctgtagttggaacctcggccaacggagctccgcagtggcagagttactccgcaccacaactaacaactgcgagtgcccgccaattggcggaacaggagtctagctcggatagctatacatccacctcgaccaccacgaccactacttcggaggagtatcagcgaatgtacgcagcccaggtgcaggcctatcaaatgcaggagcaatctggctcagagttcgattatcaggtggattacgccagtacccaggattctgtacaggactatccgtccggcaggagaagtgcccaagagtgcgtggactccctagctgtgcccctaagcacctacaagctggtcgatatggtaagggaggttacacccagtcccgtgaccactccaactcaaactcctgctcctgctgctcctacgacccgtcgcgttgtgttcaacgatgagcctgagattaaggagttaccccaactacccgcggaactagagaccataccggaagcctccgaagctgtagaagatcgcgaaggtcttgtaatcgaacagcgatgtcagattctggagagtgaacgcaagttccagcccacacccgagatcaagattgagattgccccagtgcgccaaatacctccgaccaagattcccaacccaatgcccaaggagtggattaatcccatgattcgagtcttgaccacagccccggaagttcccttccatctggtggagtgtccttttccccggccctgtggtgatgattttgaagcggaggcagccgccgccgaagcggccaaaactcaagaggttccggaacctcttcctccacaagtttctgctgctccaccggcaacagtttccgttgagccatcacctgctcccttgcgggaatctcctccccgcggatcccgactcagccaagccatggttactgctcccgagttcgagctcaagttcgcccctcccgctgaccagggcatcccacttccagaggagaccgagccctatatgccaccacccattgacacgaaaccctatttgagggaggattaccgacccaaatcaccatttgtgagtgctctaaccaccgctcccgatcgtccctttgaaggtcactttgataaagatgtgcccatccacatgattgacctgcccacccccaaagagcacctgagcatgtgtgatgccctttgcaccgccccagaacgtggttacactcccctgaatcccgagaatgctatgcatcgcgtagacgaggagcaaaagcaacaggaactcaagaagcgtgaatttcaggtgctggatcacgaggaagagctgggaatccgtccggagcctccacagtctgtcgagtactacgaaacgcggagagatcagccacggaaatcctccgcctttgcagccatgcaagcattccagccatcccgtgaacctttgtcgtcgaacacggtttcgaatgctggaagtgtggccgataccccaagagcctcgattgtgtctgccttgaaggaggaaaccgatctggagtaccagaagtatctcaaggcccagcagcgcaaccagaaaagattggactacttccaccagaaagaggaggagctctcgggtctgcagggccaacagctaacccaacttcagagggagctctcgaaccagcaacagaatcttctgagccaacagcaactgcagcaatccaagttgttgcaactgcagcagtgcgtccagagccaagagttgcagcaacaggtgcagcatctcacccagaaatcacaacagcaacctcctcaagctaaccaacagcagcaacaacagcaacagcaacggggtacccaacagcagcaacactcccaagtaacccaaagaacccaacagcaacagcaacaagtgccccaacaagtaacccaacagcaacaacaagaacactctctgctatcgcaaaccacactcgctgagacccaaacccttcaggccaatgcccagtctcagagttccgcttcctacagctccaaagcgactgcttgctctaactcctcttccacagtcccacctgccaacacctctaccgctttcgcacctgctccagctccagcacccaccagcatacctgtccgcc catcagctatcgctgtacaaagtagctactgcagcagccagttcgatgtccacgaactgatcgaggagaccgccgaggagctcgagcactcggaggtcctgttcccgccgccctccccgctgagccacctgaccaaacagggcaaagccgtacagtccggcctccacaaggcggacagcatccccaaataccagcgcaactggacggtgctacctacccagagtcccattcgcactccggaaccgcaggagctgcgcgagaacgtaccgctggcattcgtggatgctccgaaagcaccagttaccagtgactcttccactgtacatagacccattgcccaggttgctgcgccgacaactgtggttgctccttcccgggaacgggagaaggagcggcggccccagctgtcggtgcccattattgttgaggatcgatcgggtccagtaacgatggctttccaaccgttagacgaactggtgcgaccggatcaggccctgacgcccaccaggccgtacaccccgtcgctgaccaacaagccggctccaattgtgcccttctaccagacggaggagaagcttgtcttcgaggagtgctcggctacccatgccaggaactacaacgaattgaacgcctcgccttttccagacagaacacgttctccggctccgggaccaccgccaaatcccctgaatgccattcgagcaccgagaatgaaggaaccggaaaccaagtcgaatattctgtcagtttctggaggtcctcgcttgcagacgggctcaataaccactggacagagttaccagggacaacttttggctcactccgagcagagttcccagtcggccagtcagagctataaccagcaaccggagagaattacggaacaaagggtgggcaacctgaacatccaacagagggagcagtcatctcagctgcagcagcaagctcaatcgcagactcagagtcagacacgcagccaggtgggaaatactcaaatcgaaagacgtcgcaaggtcaccgaggaattcgaacgtacccagagtgctaaaactattgagatccgaactggctcccagtctgtgagtcaatcaaaggcccagtcgcagtccatcagccaggcacagacgcaggctcaatcccagtcccagaatcagtcggacacagaacgtcgctcttcgtacggtaagacaggattcgtggccagtcaggcaaagcgtctgtcctgcatggaggaagagattagcagtctgaccagccaatcgcaggctattagtgcccgggcctctgctctcggagagggctgctttcccaacctgagatcgcccacctttgactcgaagtttccacttaagccggctcccgcggagtctatagttccggggtatgcaactgttccggccgccacaaagatgctaacggcaccaccaccgggtttcctgcagcagcagcagcaacagcagcaaaggtctgccttctcgggctaccaagccacaacttcatcggtgcagcagagctcttttgcgagcagctcaaaagccacaacctcatcgctctcatcctcatcagcatctgcttcagcatcagcatccgtcgcgagatcgtcgcaaagtctaacccaagcttctgctattactaccaccactaataaccaggccaccacggcctacaggagcagcaatggcagcattaccaagcctaatctggcctcgcggccatccatcgcttccatcacagctccaggatcagcaagtgctcccgctcctgttccatcggcagctccaaccaaagctactgctccattcaaagctccgattgttccaaaatcggtgatagcgaacgccgttaacgccgctgctccgcctgcgcccgctgtctttccgccagacctgagcgatttgaacttgaactctaatgtggataattccccaggtgccggaggaaagagcgctggcgcctttggagccacctcggcgcccaagaggggcaggggtatcctgaataaggcagccggacccggagtgcgcatcccactgtgcaacagctgcaatgtgcagatcagaggaccctttatcacggcattgggccgcatctggtgcccggatcatttcatctgcgtgaacggcaactgccgtcgtccgctgcaggacattggattcgttgaggagaagggcgatctgtactgcgagtactgtttcgagaagtacctggcgcccacttgcagcaagtgcgctggcaagatcaagggtgactgtttgaatgccattggcaaacacttccatccggagtgcttcacctgcggccagtgcggcaagatctttggcaacaggcccttcttcctggaggatggaaacgcgtactgcgaggccgattggaacgagttgttcaccaccaagtgcttcgcctgcggcttccccgtggaagctggcgacagatgggtggaggccttgaaccacaactaccatagccaatgcttcaactgcacgttctgcaaacagaacctggagggtcagagcttctacaacaagggcggacgtcccttctgcaagaatcacgcgcgctaagggcgcgccatgc

Zasp52-PF PDZ + ZM (FseI and AscI sites introduced) residues 1 – 216): atggcccaaccacagctgctgcaaatcaaattgtcacgtttcgatgcccaaccctggggattccgccttcaggggggcacggacttcgctcagcccctgctggtgcaaaaggtgaacgccggcagcttgtccgagcaggctggcctccagcccggcgatgcggtggtcaagatcaatgacgtggatgtcttcaatctgcgtcacaaggatgcccaggacattgtggtgcgctccggcaacaactttgtcatcacagtgcagcgcggtggctccacctggcgcccgcatgtgacaccaactggcaatgtgccgcagcccaactcgccgtatctgcagacggtgacgaagacctctctggctcacaaacaacaggacagccagcacatcggctgtggctacaacaacgcggcccgtcccttctccaacggcggcgatggcggcgtgaagagcattgtcaataaacaatacaacaccccggttggcatttacagcgatgaatctattgcggaaacactctcggcccaggcggaggttttggctggcggtgtgctcggcgtcaacttcaagaagaacgagaaggaataccagggcgatcgctccgaggttctgaagttcctgcgcgaggaggagaccggccagtccac tcca

### F-Actin binding assay

To investigate protein binding to filamentous actin, an F-actin pelleting assay was employed. To this end, rabbit muscle actin (Akl99, Cytoskeleton) was prepared according to manufacturer’s instruction and Ca^2+^-actin exchanged to Mg^2+^-actin with exchange buffer (0.2 mM EGTA, 0.02 mM MgCl2 final) in G-buffer (5 mM Tris-HCl (pH 8.0), 0.2 mM ATP, 0.1 mM CaCl2, 0.5 mM DTT) on ice for 10 min. Mg^2+^-G-actin stock solution was diluted to 8 µM, 6 µM, 4 µM and 2 µM and co-incubated with Zasp52-ABM fragments at 8 µM in 1x polymerisation buffer (50 mM KCl, 1 mM MgCl, 1 mM EGTA, Imidazole-HCl, pH 7.0) for 1 h at RT in ultracentrifugation tubes. The samples were then centrifuged at 200,000x*g* for 1 h at 4°C. 90% of supernatant was decanted and the pellet was resuspended in the same volume of sample buffer. Supernatants and pellets were analysed via SDS PAGE and stained with Instant Blue and colorimetric image recorded with ChemiDoc XRS+ (Bio-Rad). Band intensity was quantified by densitometric scanning using Image Lab software, background intensity subtracted and ABM fraction binding quantified (Int[ABMbound]/Int[ABMbound]+Int[ABMunbound]).

### Immunoprecipitation of GFP-labelled proteins from Drosophila embryos

To identify proteins associated with Zasp52 in epithelia, co-immunoprecipitations against GFP were performed using either the *Zasp52-GFP[ZCL423]* or *Zasp52-GFP[G00189]* endogenous protein trap lines as well as wild-type *yw* flies (Morin et al., 2001). *Armadillo-YFP[CPTI001198]* was used as a control for co-immunoprecipitation of a junctional cytoplasmic protein. To prevent the contamination of the sample with muscle tissue, the collection protocol consisted of a one hour pre-lay collection to eliminate embryos retained in the mother (discarded), and was then followed by a 9-hour collection, resulting in embryos between 0 and 9 hpf and thus prior to muscle development which commences at 9:20 hpf. Dechorionated, frozen embryos were placed on ice and immediately covered with 100 µl chilled lysis buffer (50 mM Tris pH 7.4, 150 mM KCl, 0.5 mM EDTA, 0.1 % Glycerol, 0.01 % Triton X-100, 1200 μg/ml benzamidine, 40 μg/ml chymostatin, 40 μg/ml antipain, 2 μg/ml leupeptin, 0.96 μg/ml pefabloc, 0.5 mM PMSF). Each sample contained roughly the same mass of embryos, as different samples were used per experiment in order to observe differences between sample types, e.g. wild-type versus Zasp52-GFP, and quantitative analysis of triplicate samples used. If necessary, different collections of equivalent laying conditions were pooled to create one sample. Per sample, 500 µl of lysis buffer were added and total sample transferred into a glass dounce-homogenizer. Embryos were lysed with 20 strokes and a subsequent 30 min incubation on ice. To separate the protein-containing aqueous phase from lipids and cell debris, samples were centrifuged at max speed in a table top centrifuge for 10 min at 4°C. The aqueous phase was carefully collected and added to equal amounts of GFP-nanobody coated magnetic beads (Chromotek), equilibrated in lysis buffer. Samples were incubated on a rotating shaker for 1.5 h at 4°C. Samples were then washed three times with wash buffer (10 mM Tris pH 7.4, 150 mM KCl, 0.5 mM EDTA, 0.1 % Glycerol, 0.5 mM PMSF) and then either eluted in sample buffer for subsequent analysis by western blotting, or the wash buffer decanted, beads resuspended in 50 mM ammonium bicarbonate and stored at -20°C prior to mass spectrometry analysis.

### Mass-spectrometry processing of immunoprecipitation samples

Proteins were prepared for enzymatic cleavage from magnetic beads by submersion in 50 mM ammonium hydrogen carbonate, pH 8.0. The solution containing bead-bound protein was digested with 0.5 µg of trypsin for 60 min at 37 °C in a thermomixer, shaking at 800 rpm. This was followed by another overnight digestion step for which an additional 1µg trypsin was added and the samples digested under the same conditions as above. The reaction was terminated by adding formic acid to a final concentration of 2% v/v. The output of the digestion reaction was analysed using a nano-scale capillary LC-MS/MS in a Ultimate U3000 HPLC setup (ThermoScientific Dionex, San Jose, USA) to deliver a flow of approximately 300 nl/min. A C18 Acclaim PepMap100 5 µm, 100 µm x 20 mm nanoViper (ThermoScientific Dionex, San Jose, USA), trapped the peptides before separation on a C18 Acclaim PepMap100 3 µm, 75 µm x 150 mm nanoViper (ThermoScientific Dionex, San Jose, USA). The Peptides were eluted from the column via an acetonitrile gradient. The analytical column outlet was immediately interfaced by a modified nano-flow electrospray ionisation source, coupled to a hybrid dual pressure linear ion trap mass spectrometer (Orbitrap Velos, ThermoScientific, San Jose, USA). The data acquisition was carried out, using a resolution of 30,000 for the full MS spectrum, then followed by ten MS/MS spectra in the linear ion trap. All MS spectra were collected over a m/z range of 300–2000 and MS/MS scans were collected with a threshold energy of 35 for collision induced dissociation. All raw files were processed with MaxQuant 1.5.5.1 (Cox and Mann, 2008) using standard settings and searched against the UniProt KB with the Andromeda search engine (Cox et al., 2011) integrated into the MaxQuant software suite. Enzyme search specificity was Trypsin/P for both endoproteinases. One or two missed cleavages for each peptide were allowed. Carbamido-methylation of cysteines as selected as fixed modification with oxidized methionine and protein N-acetylation set as variable modifications. The search was conducted with an initial mass tolerance of 6 ppm for the precursor ion and 0.5 Da for MS/MS spectra. The false discovery rate was fixed at 1% at the peptide and protein level.

### Quantitative analysis of mass-spectrometric data of co-immunoprecipitations in Perseus

Downstream statistical analysis was carried out using the Perseus interface of MaxQuant (https://maxquant.net/perseus/). Prior to statistical analysis, peptides mapped to known contaminants, e.g. human Keratin, reverse sequence hits and protein groups only identified by site were removed. Only protein groups identified with at least two peptides, of which one had to be unique, and two quantitation events were considered for data analysis with Perseus. Experiments analysed were 3 for Zasp52-GFP and Armadillo-YFP each and 2 for wild type. To compare samples of interest to control, the triplicate or duplicate data sets were transformed to Log2(x), missing values replaced by imputation and the volcano plot tool used to compare two data sets in which the x-axis represents a difference score and the y-axis visualises the negative, logarithmic p value generated by Student T-test. The significance threshold was set to a p value of 0.75, so that known interaction partners in the *Armadillo-YFP* control sample were identified.

### Computational modelling

We represent an epithelial tissue by a 2D vertex model of 127 hexagonal cells initially arranged in a hexagonal orientation, essentially following previous (Durney and Feng, 2021; Durney et al., 2018). In brief, an individual cell has six peripheral nodes and one central node connected by passively elastic edges. To represent a supracellular actomyosin cable that is enriched by myosin across several junctional boundaries, we have taken select edges (indicated in bold black, Fig. 7A, B) and endowed them with two different properties: an increased elastic modulus, μ, and a resistance to bending. The latter is implemented by a penalty against deviation in the angle of adjacent edges resulting in a restoring force, *F_b_*. Select edges (indicated in green, Fig. 7A) are prescribed to contract and generate active pulling forces. This contraction pulls on the tissue exterior of the boundary compressing the interior region.

The total force, ***F****_i_*, on node *i* is given by,

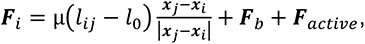

where *l_ij_* is the current edge length between nodes *i* and *j*, *l*_0_ is the edge rest length, ***F****_b_* is the force due to bending (only on edges associated with the supracellular actomyosin cable) and ***F****_active_* is the active force that causes contraction of select edges. Nodal motion is governed by over-damped dynamics with a viscous friction factor *η*,

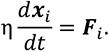

The force, ***F****_b_* = *k*(θ – π), tends to align the adjacent edges. As the tissue is in an initial hexagonal configuration, this term circularizes the supracellular cable and reduces its circumference. To maintain a zero-energy ground state, we set the bending modulus, *k*, and then allow the tissue to relax to equilibrium. For edges of the supracellular cable, we adopt the equilibrium length to be its new rest length. This procedure is repeated until there are no more changes in *l_ij_* Ɏ *i*,*j*. This guarantees a consistent ground state as μ is increased.

Tissue deformation is initiated by *F_active_* contracting specific edges.

The main output of the model is the average radial movement of the *n* peripheral nodes of the epithelial tissue,

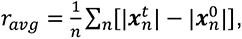

where the superscript denotes simulation time.

## Acknowledgements

The authors would like to thank the following people; for fly stocks: Frieder Schöck, Debbie Andrew, Magali Suzanne, Isabel Palacios; for reagents: Emmanuel Derivery; We thank members of the lab and Cell Biology division for input on the manuscript. K.R., D.J.A., V.J.P.H., and T.S. were supported by the Medical Research Council (file reference number U105178780), C.D. and J.F by Natural Sciences and Engineering Research Council Canada (2019-04162). This work was supported by the Medical Research Council, as part of United Kingdom Research and Innovation (also known as UK Research and Innovation) [MRC file reference number U105178780]. For the purpose of open access, the author has applied a CC BY public copyright licence to any Author Accepted Manuscript version arising

## Author contribution

Conceptualisation, K.R. and D.J.A.; Methodology, K.R., D.J.A., V.J.P.H, T.S., C.D and J.F.; Investigation, K.R., D.J.A., C.D.; Writing-Original Draft, K.R., D.J.A.; Funding Acquisition, K.R., J.F.

## Declaration of Interests

The authors declare no competing interests.

***Supplemental Movie S1, related to Figure 7. Analysis of cell vertex movement inside and outside the salivary gland placode boundary.***

Time lapse movie of a *sqhGPF; UbiRFP* embryo (myosin highlighted by SqhGFP and cell membranes by UbiRFP) showing the salivary gland placode and surrounding epidermal cells starting from late stage 10 onwards. Time interval between frames is 1min:30sec.

***Supplemental Table S1, related to Figure 3. Proteins identified by mass-spectrometric analysis upon anti-GFP co-immunoprecipitation from Zasp52-GFP-Z embryos in comparison to wt embryos.***

All hits (interactors) identified by the mass-spectrometric analysis, listing Difference and -log(P value), as well as protein IDs and actual gene names as used in *Drosophila melanogaster*. All hits are also manually classified as being involved in cell adhesion, microtubule cytoskeleton, actin cytoskeleton, translation, transcription, muscle development and function or mitochondrial function.

***Supplemental Table S2, related to Figure 3. Proteins identified by mass-spectrometric analysis upon anti-GFP co-immunoprecipitation from Armadillo-YFP embryos in comparison to wt embryos.***

All hits (interactors) identified by the mass-spectrometric analysis, listing Difference and -log(P value), as well as protein IDs and actual gene names as used in *Drosophila melanogaster*. All hits are also manually classified as being involved in cell adhesion, microtubule cytoskeleton, actin cytoskeleton, translation, transcription, muscle development and function or mitochondrial function.

---

**Supplemental Figure S1, related to Figures 1, 2, 4, 5 and 6.**
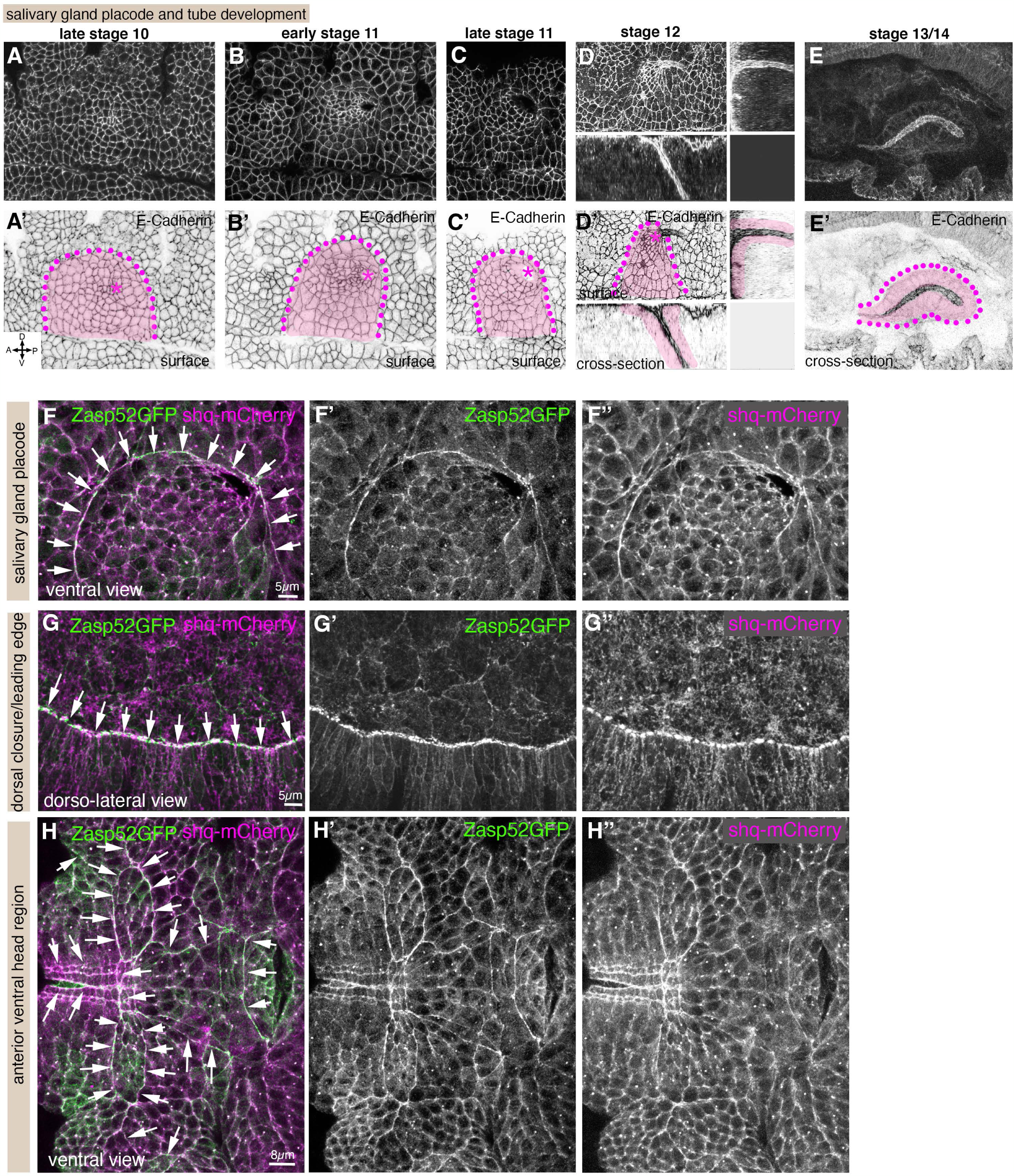
Embryonic development of the salivary gland and Zasp52 localisation in developmental actomyosin cables. **A-E ’** Development of the salivary gland begins at late stage 10 from a flat epidermal placode of about one hundred cells each on either side of the ventral midline (**A, A’**). **B-C’** Cells invaginate through a focal point in the dorsal-posterior corner. **D-E’** A narrow-lumen tube is formed as soon as cells internalise and the tube extends internally with more cells invaginating. Purple dotted lines mark the boundary of the placode, pink overlay indicates the salivary gland cells, and asterisks mark the position of the invagination point or pit. Surface views (**A-C’** and main panels in **D**, **D’**) and cross section views (small panels in **D**, **D’**; and **E**, **E’**) are shown. E-Cadherin labelling of apical junctional cell outlines is shown, with lower panels showing inverse label to better visualise labels. **F-H’’** Zasp52 localisation in actomyosin cables during embryonic development in comparison to Sqh (non-muscle myosin II regulatory light chain). Zasp52-GFP-Z is in green, Sqh-mCherry in magenta. **F-F’’** shows the cable at the boundary of the salivary gland placode, **G-G’’** the cable during dorsal closure at the leading edge-amnioserosa interface, **H-H’’** the network of anterior ventral cables during head involution. Arrows point to the position of the cables.

**Supplemental Figure S2, related to Figure 1.**
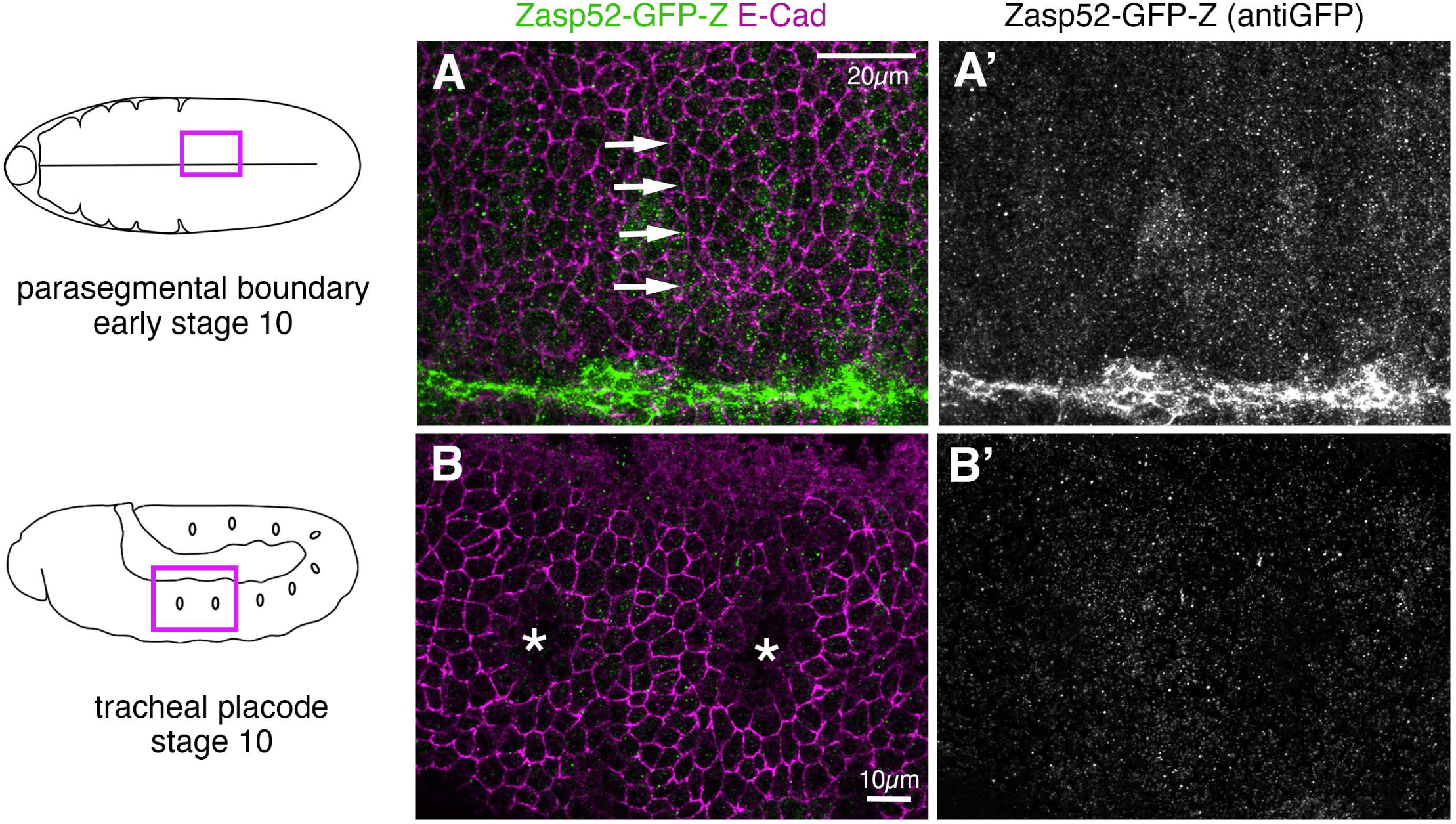
Zasp52-GFP is not found in parasegmental or tracheal pit actomyosin cables. **A**, **A’** A parasegmental boundary within the embryonic epidermis at early stage 10. Note that Zasp52-GFP in green in **A** and as a single channel in **A’** is strongly expressed in the ventral midline but not at all enriched at parasegmental boundaries (arrows in **A**) at this stage. **B**, **B’** Two tracheal placodes within the embryonic epidermis at stage 10, the invagination points are marked by asterisks. Zasp52-GFP is not enriched in any junctions within these placodes.

**Supplemental Figure S3, related to Figure 2.**
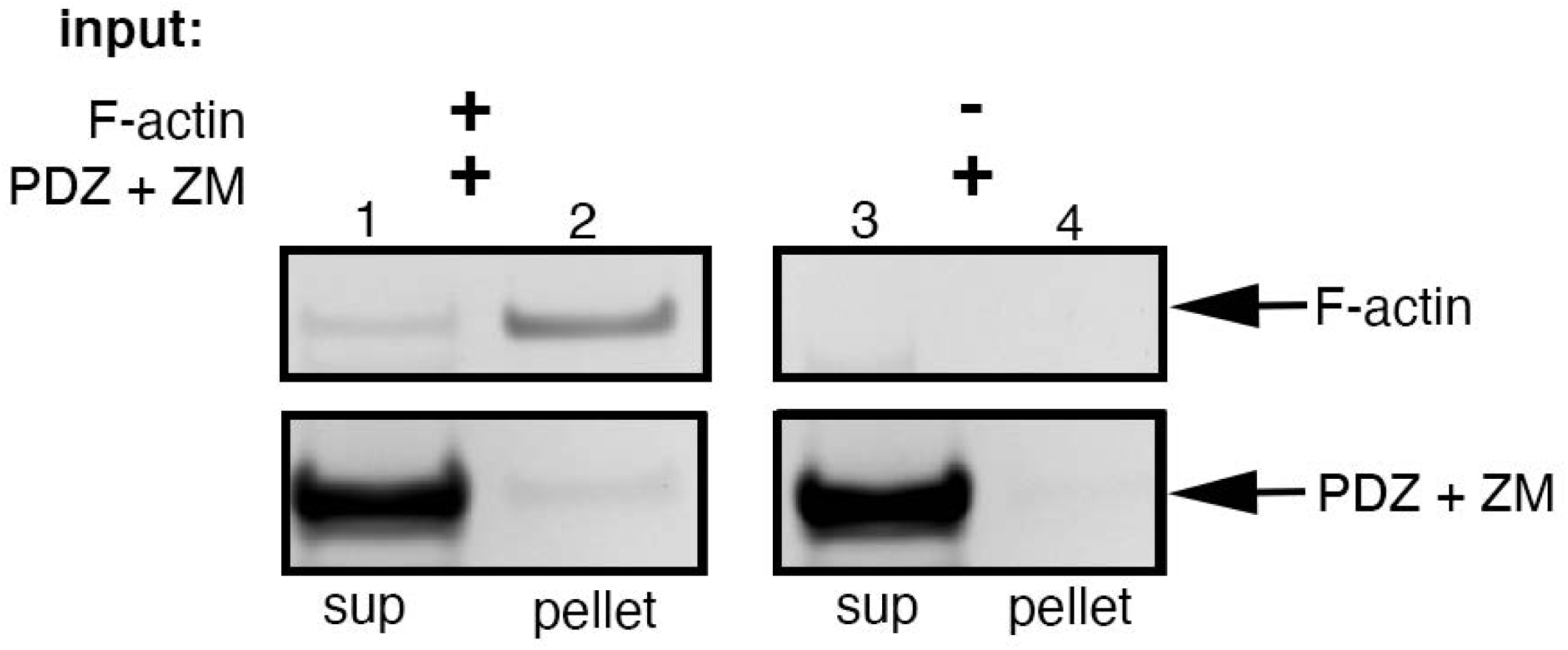
F-actin binding of Zasp52 N-terminus. The Zasp52 PDZ domain and Zasp motif were expressed in bacteria and purified protein was used in an F-actin pelleting assay. F-actin can co-sediment a fraction of PDZ+ZM (lane 2), whereas PDZ+ZM alone does not pellet (lane 4) but remains in the supernatant (lane 3).

**Supplemental Figure S4, related to Figure 3.**
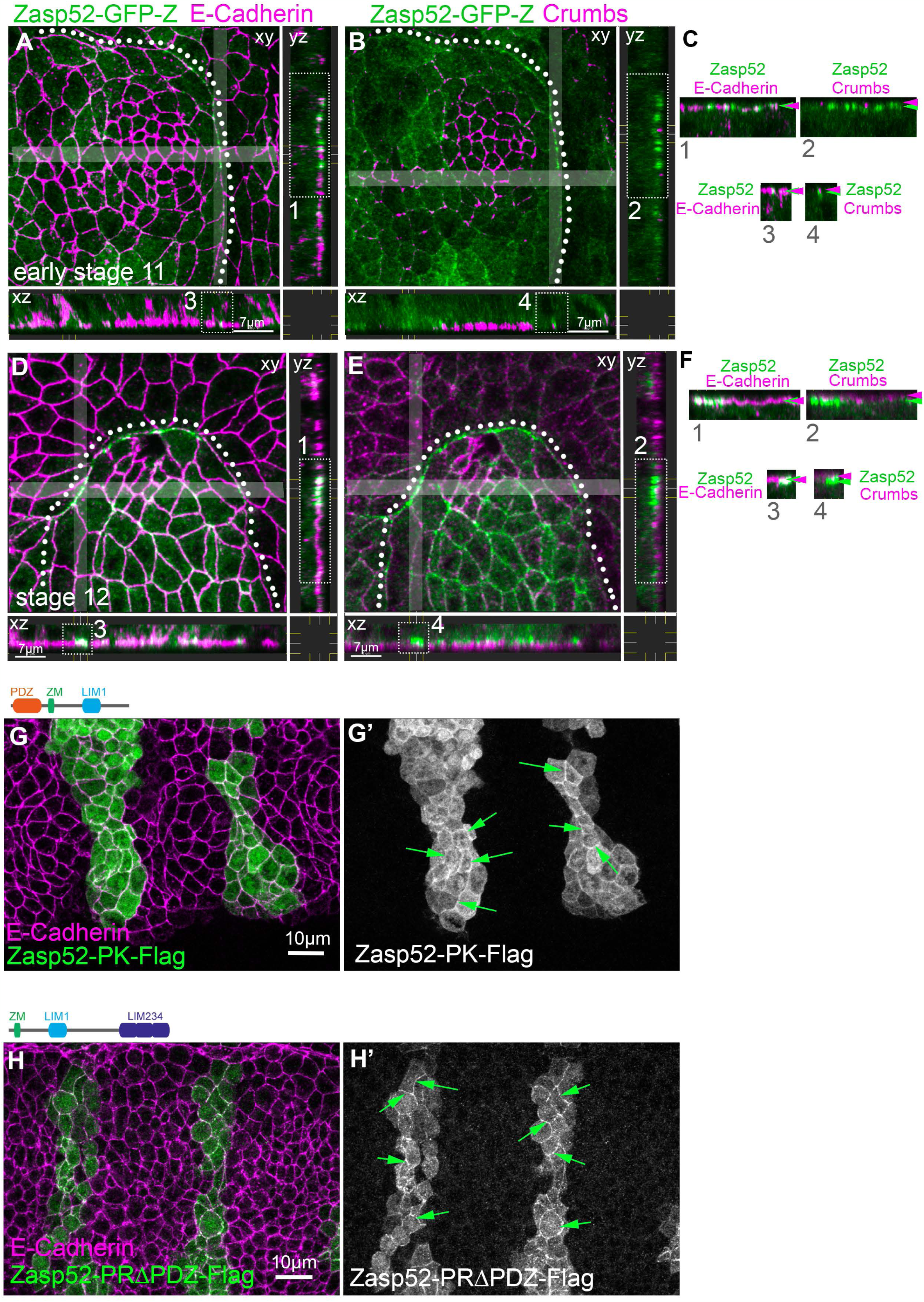
Endogenous and ectopic Zasp52 localisation at cell-cell junctions. **A-F** Localisation of Zasp52GFP in actomyosin cables along the apical-lateral side of cells. Examples of localisation in the actomyosin cable surrounding the salivary gland placode at early stage 11 (**A**-**C**) and at stage 12 (**D**-**F**). Zasp52-GFP (green) localisation is compared to E-Cadherin at adherens junctions (magenta in **A**, **D**) and to Crumbs in the marginal zone (magenta in **B**, **E**). Surface stack views (xy) with corresponding cross sections (xz and yz) are shown, with the width of the section displayed indicated by grey shading. White dotted boxes in the cross sections (labelled 1-4) are displayed again in **C** and **D** to compare the localisation of Zasp52-GFP, E-Cadherin and Crumbs along the apical to basal direction, apical is up in **C** and **D**. Coloured arrowheads indicate the position of Zasp52GFP (green) or E-Cadherin and Crumbs (magenta) as indicated. White dotted lines indicate the boundary of the salivary gland placodes shown. Scale bars are 7µm. **G**, **G’** Stripe-overexpression of the Zasp52-PK isoform (as *UAS-Zasp52-PK-Flag*), containing the PDZ domain, Zasp motif and LIM1 domain, under control of *enGal4* leads to it being localised to cell-cell junctions in addition to a cytoplasmic pool. **H**, **H’** Stripe-overexpression of the Zasp52-PRΔPDZ isoform (as *UAS-Zasp52-PRΔPDZ - Flag*), containing the ZM motif and LIM 1-4 domains, under control of *enGal4* leads to it being localised to cell-cell junctions in addition to a cytoplasmic pool. Zasp52 isoforms are in green and as a single channel in **G’**, **H’** and E-Cadherin to label cell junctions in magenta. Arrows point to junctional localisation of the ectopically expressed Zasp52 protein variants.

**Supplemental Figure S5, related to Figure 6.**
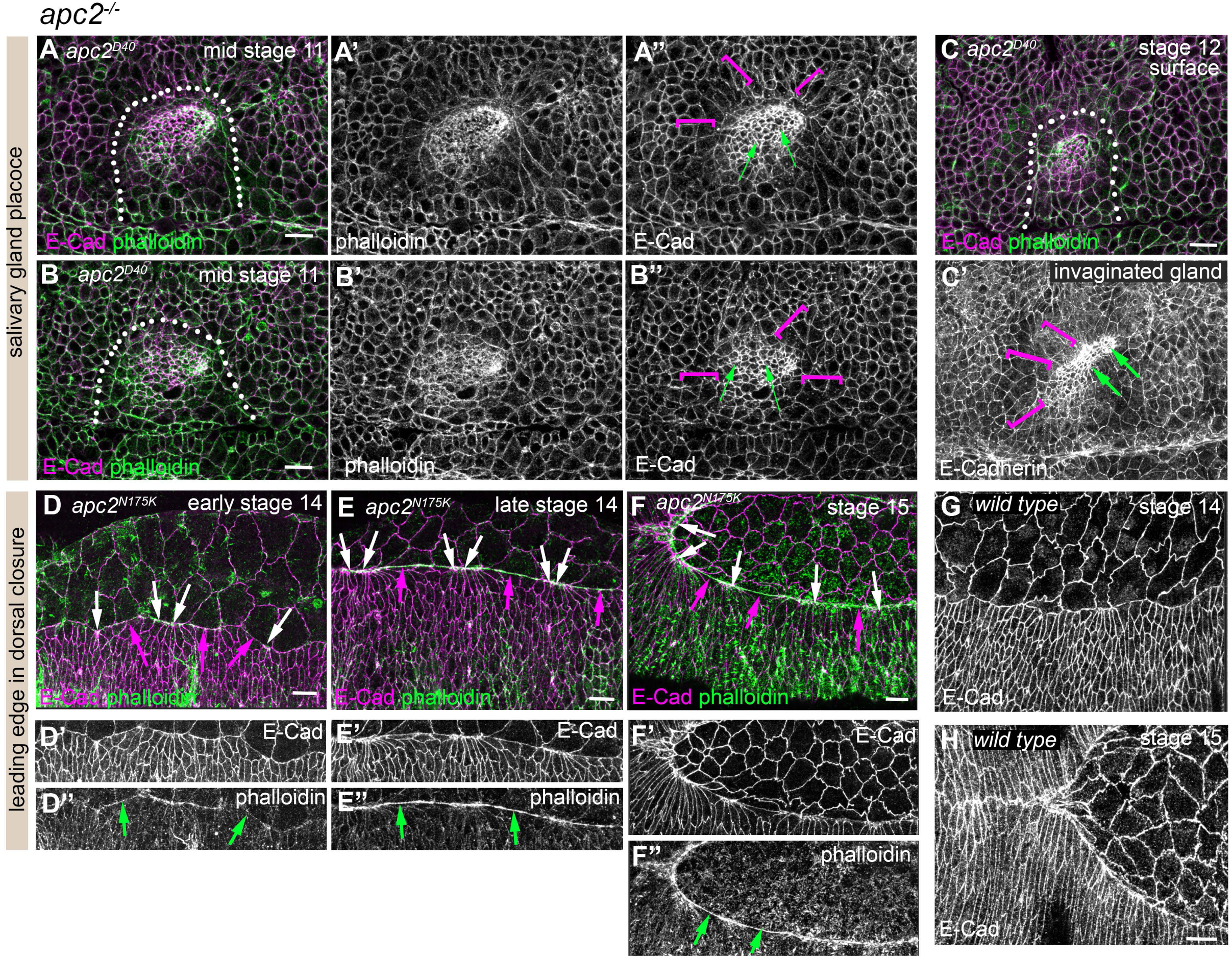
apc2^-/-^ mutant embryonic phenotypes. **A-B’** Two examples of *apc^D40^* mutant salivary gland placodes, at mid stage 11 when invagination has begun. Note that secretory cells of the placode appear to be more constricted than usual (green arrows; see Fig.6C for comparison to control), and cells immediately surrounding the placode boundary are overstretched (magenta brackets). **C**,**C’** The invaginated portion of the salivary gland at stage 12 in an *apc^D40^* mutant embryo shows a too wide lumen, **C** shows the surface views and **C’** a projection of the surface and invaginated portion. **D-H** The leading edge during dorsal closure in *apc2^N175K^* mutant embryos is aberrant, with failed actin accumulation in some areas from stage 14-15, and with some overconstricted and some too relaxed cells at the leading edge itself (**D**-**F’’**; green arrows point to low cortical actin, white arrows to overconstricted cells and magenta arrows to overstretched cells). **G** and **H** show wild-type leading edges at stage 14 and 15, respectively. In all colour panels E-Cadherin is shown in magenta and phalloidin to label F-actin is shown in green, and staining is indicated on single channel panels.

## References

1. Ahmed, Y., Nouri, A., and Wieschaus, E. (2002). Drosophila Apc1 and Apc2 regulate Wingless transduction throughout development. Development 129, 1751–1762. 10.1242/dev.129.7.1751.

2. Blanchard, G.B., Kabla, A.J., Schultz, N.L., Butler, L.C., Sanson, B., Gorfinkiel, N., Mahadevan, L., and Adams, R.J. (2009). Tissue tectonics: morphogenetic strain rates, cell shape change and intercalation. Nat Methods 6, 458–464.

3. Choi, W., Jung, K.C., Nelson, K.S., Bhat, M.A., Beitel, G.J., Peifer, M., and Fanning, A.S. (2011). The single Drosophila ZO-1 protein Polychaetoid regulates embryonic morphogenesis in coordination with Canoe/afadin and Enabled. Mol Biol Cell 22, 2010–2030. 10.1091/mbc.E10-12-1014.

4. Chou, T.B., and Perrimon, N. (1992). Use of a yeast site-specific recombinase to produce female germline chimeras in Drosophila. Genetics 131, 643–653. 10.1093/genetics/131.3.643.

5. Chugh, P., and Paluch, E.K. (2018). The actin cortex at a glance. J Cell Sci 131. 10.1242/jcs.186254.

6. Cox, J., and Mann, M. (2008). MaxQuant enables high peptide identification rates, individualized p.p.b.-range mass accuracies and proteome-wide protein quantification. Nature biotechnology 26, 1367–1372. 10.1038/nbt.1511.

7. Cox, J., Neuhauser, N., Michalski, A., Scheltema, R.A., Olsen, J.V., and Mann, M. (2011). Andromeda: a peptide search engine integrated into the MaxQuant environment. J Proteome Res 10, 1794–1805. 10.1021/pr101065j.

8. Desai, R., Sarpal, R., Ishiyama, N., Pellikka, M., Ikura, M., and Tepass, U. (2013). Monomeric alpha-catenin links cadherin to the actin cytoskeleton. Nat Cell Biol 15, 261–273. 10.1038/ncb2685.

9. Ducuing, A., Keeley, C., Mollereau, B., and Vincent, S. (2015). A DPP-mediated feed-forward loop canalizes morphogenesis during Drosophila dorsal closure. J Cell Biol 208, 239–248. 10.1083/jcb.201410042.

10. Ducuing, A., and Vincent, S. (2016). The actin cable is dispensable in directing dorsal closure dynamics but neutralizes mechanical stress to prevent scarring in the Drosophila embryo. Nat Cell Biol 18, 1149–1160. 10.1038/ncb3421.

11. Durney, C.H., and Feng, J.J. (2021). A three-dimensional vertex model forDrosophilasalivary gland invagination. Phys Biol 18. 10.1088/1478-3975/abfa69.

12. Durney, C.H., Harris, T.J.C., and Feng, J.J. (2018). Dynamics of PAR Proteins Explain the Oscillation and Ratcheting Mechanisms in Dorsal Closure. Biophys J 115, 2230–2241. 10.1016/j.bpj.2018.10.014.

13. Ebrahim, S., Fujita, T., Millis, B.A., Kozin, E., Ma, X., Kawamoto, S., Baird, M.A., Davidson, M., Yonemura, S., Hisa, Y., et al. (2013). NMII forms a contractile transcellular sarcomeric network to regulate apical cell junctions and tissue geometry. Curr Biol 23, 731–736. 10.1016/j.cub.2013.03.039.

14. Fernandez-Gonzalez, R., Simoes Sde, M., Roper, J.C., Eaton, S., and Zallen, J.A. (2009). Myosin II dynamics are regulated by tension in intercalating cells. Dev Cell 17, 736–743. S1534-5807(09)00385-2 [pii]. 10.1016/j.devcel.2009.09.003.

15. Finegan, T.M., Hervieux, N., Nestor-Bergmann, A., Fletcher, A.G., Blanchard, G.B., and Sanson, B. (2019). The tricellular vertex-specific adhesion molecule Sidekick facilitates polarised cell intercalation during Drosophila axis extension. PLoS biology 17, e3000522. 10.1371/journal.pbio.3000522.

16. Furukawa, K.T., Yamashita, K., Sakurai, N., and Ohno, S. (2017). The Epithelial Circumferential Actin Belt Regulates YAP/TAZ through Nucleocytoplasmic Shuttling of Merlin. Cell reports 20, 1435–1447. 10.1016/j.celrep.2017.07.032.

17. Galea, G.L., Cho, Y.J., Galea, G., Mole, M.A., Rolo, A., Savery, D., Moulding, D., Culshaw, L.H., Nikolopoulou, E., Greene, N.D.E., and Copp, A.J. (2017). Biomechanical coupling facilitates spinal neural tube closure in mouse embryos. Proc Natl Acad Sci U S A 114, E5177–E5186. 10.1073/pnas.1700934114.

18. Girdler, G.C., and Röper, K. (2014). Controlling cell shape changes during salivary gland tube formation in Drosophila. Semin Cell Dev Biol 31, 74–81. 10.1016/j.semcdb.2014.03.020.

19. Gonzalez-Morales, N., Xiao, Y.S., Schilling, M.A., Marescal, O., Liao, K.A., and Schöck, F. (2019). Myofibril diameter is set by a finely tuned mechanism of protein oligomerization in Drosophila. eLife 8. 10.7554/eLife.50496.

20. Guirao, B., Rigaud, S.U., Bosveld, F., Bailles, A., Lopez-Gay, J., Ishihara, S., Sugimura, K., Graner, F., and Bellaiche, Y. (2015). Unified quantitative characterization of epithelial tissue development. eLife 4. 10.7554/eLife.08519.

21. Hamada, F., and Bienz, M. (2002). A Drosophila APC tumour suppressor homologue functions in cellular adhesion. Nat Cell Biol 4, 208–213. 10.1038/ncb755.

22. Hanson, J., Yang, Y., Paliwal, K., and Zhou, Y. (2017). Improving protein disorder prediction by deep bidirectional long short-term memory recurrent neural networks. Bioinformatics 33, 685–692. 10.1093/bioinformatics/btw678.

23. Hashimoto, H., and Munro, E. (2019). Differential Expression of a Classic Cadherin Directs Tissue-Level Contractile Asymmetry during Neural Tube Closure. Dev Cell 51, 158–172 e154. 10.1016/j.devcel.2019.10.001.

24. Herrera-Perez, R.M., and Kasza, K.E. (2018). Biophysical control of the cell rearrangements and cell shape changes that build epithelial tissues. Curr Opin Genet Dev 51, 88–95. 10.1016/j.gde.2018.07.005.

25. Hug, C., Miller, T.M., Torres, M.A., Casella, J.F., and Cooper, J.A. (1992). Identification and characterization of an actin-binding site of CapZ. J Cell Biol 116, 923–931. 10.1083/jcb.116.4.923.

26. Jacinto, A., Wood, W., Woolner, S., Hiley, C., Turner, L., Wilson, C., Martinez-Arias, A., and Martin, P. (2002). Dynamic analysis of actin cable function during Drosophila dorsal closure. Curr Biol 12, 1245–1250.

27. Jani, K., and Schöck, F. (2007). Zasp is required for the assembly of functional integrin adhesion sites. J Cell Biol 179, 1583–1597. 10.1083/jcb.200707045.

28. Jones, D.T., and Cozzetto, D. (2015). DISOPRED3: precise disordered region predictions with annotated protein-binding activity. Bioinformatics 31, 857–863. 10.1093/bioinformatics/btu744.

29. Kim, T., Cooper, J.A., and Sept, D. (2010). The interaction of capping protein with the barbed end of the actin filament. J Mol Biol 404, 794–802. 10.1016/j.jmb.2010.10.017.

30. Klausen, M.S., Jespersen, M.C., Nielsen, H., Jensen, K.K., Jurtz, V.I., Sonderby, C.K., Sommer, M.O.A., Winther, O., Nielsen, M., Petersen, B., and Marcatili, P. (2019). NetSurfP-2.0: Improved prediction of protein structural features by integrated deep learning. Proteins 87, 520–527. 10.1002/prot.25674.

31. Letizia, A., He, D., Astigarraga, S., Colombelli, J., Hatini, V., Llimargas, M., and Treisman, J.E. (2019). Sidekick Is a Key Component of Tricellular Adherens Junctions that Acts to Resolve Cell Rearrangements. Dev Cell 50, 313–326 e315. 10.1016/j.devcel.2019.07.007.

32. Liao, K.A., Gonzalez-Morales, N., and Schöck, F. (2020). Characterizing the actin-binding ability of Zasp52 and its contribution to myofibril assembly. PLoS One 15, e0232137. 10.1371/journal.pone.0232137.

33. Major, R.J., and Irvine, K.D. (2006). Localization and requirement for Myosin II at the dorsal-ventral compartment boundary of the Drosophila wing. Dev Dyn 235, 3051–3058. 10.1002/dvdy.20966.

34. Martin, P., and Lewis, J. (1992). Actin cables and epidermal movement in embryonic wound healing. Nature 360, 179–183. 10.1038/360179a0.

35. Mege, R.M., and Ishiyama, N. (2017). Integration of Cadherin Adhesion and Cytoskeleton at Adherens Junctions. Cold Spring Harb Perspect Biol 9. 10.1101/cshperspect.a028738.

36. Monier, B., Pelissier-Monier, A., Brand, A.H., and Sanson, B. (2010). An actomyosin-based barrier inhibits cell mixing at compartmental boundaries in Drosophila embryos. Nat Cell Biol 12, 60–65; sup pp 61-69. ncb2005 [pii]. 10.1038/ncb2005.

37. Morin, X., Daneman, R., Zavortink, M., and Chia, W. (2001). A protein trap strategy to detect GFP-tagged proteins expressed from their endogenous loci in Drosophila. Proc Natl Acad Sci U S A 98, 15050–15055. 10.1073/pnas.261408198.

38. Narita, A., Takeda, S., Yamashita, A., and Maeda, Y. (2006). Structural basis of actin filament capping at the barbed-end: a cryo-electron microscopy study. EMBO J 25, 5626–5633. 10.1038/sj.emboj.7601395.

39. Orsulic, S., and Peifer, M. (1996). An in vivo structure-function study of armadillo, the beta-catenin homologue, reveals both separate and overlapping regions of the protein required for cell adhesion and for wingless signaling. J Cell Biol 134, 1283–1300. 10.1083/jcb.134.5.1283.

40. Pare, A.C., Naik, P., Shi, J., Mirman, Z., Palmquist, K.H., and Zallen, J.A. (2019). An LRR Receptor-Teneurin System Directs Planar Polarity at Compartment Boundaries. Dev Cell 51, 208–221 e206. 10.1016/j.devcel.2019.08.003.

41. Pare, A.C., Vichas, A., Fincher, C.T., Mirman, Z., Farrell, D.L., Mainieri, A., and Zallen, J.A. (2014). A positional Toll receptor code directs convergent extension in Drosophila. Nature 515, 523–527. 10.1038/nature13953.

42. Rodriguez-Diaz, A., Toyama, Y., Abravanel, D.L., Wiemann, J.M., Wells, A.R., Tulu, U.S., Edwards, G.S., and Kiehart, D.P. (2008). Actomyosin purse strings: renewable resources that make morphogenesis robust and resilient. HFSP J 2, 220–237. 10.2976/1.2955565.

43. Röper, K. (2012). Anisotropy of Crumbs and aPKC Drives Myosin Cable Assembly during Tube Formation. Dev Cell 23, 939–953. 10.1016/j.devcel.2012.09.013 S1534-5807(12)00424-8 [pii].

44. Röper, K. (2013). Supracellular actomyosin assemblies during development. Bioarchitecture 3, 45–49. 10.4161/bioa.25339.

45. Röper, K. (2015). Integration of cell-cell adhesion and contractile actomyosin activity during morphogenesis. Curr Top Dev Biol 112, 103–127. 10.1016/bs.ctdb.2014.11.017.

46. Sanchez-Corrales, Y.E., Blanchard, G., and Röper, K. (2021). Correct regionalisation of a tissue primordium is essential for coordinated morphogenesis. BioRxiv. 2020.08.29.273219.

47. Sanchez-Corrales, Y.E., Blanchard, G.B., and Röper, K. (2018). Radially patterned cell behaviours during tube budding from an epithelium. eLife 7. 10.7554/eLife.35717.

48. Sanchez-Corrales, Y.E., and Röper, K. (2018). Alignment of cytoskeletal structures across cell boundaries generates tissue cohesion during organ formation. Curr Opin Cell Biol 55, 104–110. 10.1016/j.ceb.2018.07.001.

49. Sarpal, R., Pellikka, M., Patel, R.R., Hui, F.Y., Godt, D., and Tepass, U. (2012). Mutational analysis supports a core role for Drosophila alpha-catenin in adherens junction function. J Cell Sci 125, 233–245. jcs.096644 [pii]. 10.1242/jcs.096644.

50. Sidor, C., and Röper, K. (2016). Genetic Control of Salivary Gland Tubulogenesis in Drosophila. In Organogenetic Gene Networks, J. Castelli-Gair Hombría, and P. Bovolenta, eds. (Srpinger International Publishing), pp. 125–149. DOI 10.1007/978-3-319-42767-6_5.

51. Sidor, C., Stevens, T.J., Jin, L., Boulanger, J., and Röper, K. (2020). Rho-Kinase Planar Polarization at Tissue Boundaries Depends on Phospho-regulation of Membrane Residence Time. Dev Cell 52, 364–378 e367. 10.1016/j.devcel.2019.12.003.

52. Song, S., Eckerle, S., Onichtchouk, D., Marrs, J.A., Nitschke, R., and Driever, W. (2013). Pou5f1-dependent EGF expression controls E-cadherin endocytosis, cell adhesion, and zebrafish epiboly movements. Dev Cell 24, 486–501. 10.1016/j.devcel.2013.01.016.

53. Stronach, B. (2014). Extensive nonmuscle expression and epithelial apicobasal localization of the Drosophila ALP/Enigma family protein, Zasp52. Gene Expr Patterns 15, 67–79. 10.1016/j.gep.2014.05.002.

54. Tepass, U., Gruszynski-DeFeo, E., Haag, T.A., Omatyar, L., Torok, T., and Hartenstein, V. (1996). shotgun encodes Drosophila E-cadherin and is preferentially required during cell rearrangement in the neurectoderm and other morphogenetically active epithelia. Genes Dev 10, 672–685.

55. Tomancak, P., Beaton, A., Weiszmann, R., Kwan, E., Shu, S., Lewis, S.E., Richards, S., Ashburner, M., Hartenstein, V., Celniker, S.E., and Rubin, G.M. (2002). Systematic determination of patterns of gene expression during Drosophila embryogenesis. Genome Biol 3, RESEARCH0088. 10.1186/gb-2002-3-12-research0088.

56. Urnavicius, L., Zhang, K., Diamant, A.G., Motz, C., Schlager, M.A., Yu, M., Patel, N.A., Robinson, C.V., and Carter, A.P. (2015). The structure of the dynactin complex and its interaction with dynein. Science 347, 1441–1446. 10.1126/science.aaa4080.

57. Wood, W., Jacinto, A., Grose, R., Woolner, S., Gale, J., Wilson, C., and Martin, P. (2002). Wound healing recapitulates morphogenesis in Drosophila embryos. Nat Cell Biol 4, 907–912. 10.1038/ncb875.

58. Yamashita, A., Maeda, K., and Maeda, Y. (2003). Crystal structure of CapZ: structural basis for actin filament barbed end capping. EMBO J 22, 1529–1538. 10.1093/emboj/cdg167.

